# The metastatic capacity of high-grade serous ovarian cancer cells changes along disease progression: inhibition by mifepristone

**DOI:** 10.1101/2022.05.07.491021

**Authors:** Sabrina J. Ritch, Abu S. M. Noman, Alicia A. Goyeneche, Carlos M. Telleria

**Author notes:** Alicia A. Goyeneche and Carlos M. Telleria share senior co-authorship. Author’s e-mails: Sabrina J. Ritch, Abu S. M. Noman, Alicia A. Goyeneche, Carlos M. Telleria.

## Abstract

**Background:** Reductionist two-dimensional (2D) in vitro assays have long been the standard for studying the metastatic abilities of cancer cells. However, tri-dimensional (3D) organotypic models provide a more complex environment, closer to that seen in patients, and thereby provide a more accurate representation of their true capabilities. Our laboratory has previously shown that the antiprogestin and antiglucocorticoid mifepristone can reduce the growth, adhesion, migration, and invasion of various aggressive cancer cells assessed using 2D assays. In this study, we characterize the metastatic capabilities of high-grade serous ovarian cancer cells generated along disease progression, in both 2D and 3D assays, and the ability of cytostatic doses of mifepristone to inhibit them.

**Methods:** High-grade serous ovarian cancer cells collected from two separate patients at different stages of their disease were used throughout the study. The 2D wound healing and Boyden chamber assays were used to study migration, while a layer of extracellular matrix was added to the Boyden chamber to study invasion. A 3D organotypic model, composed of fibroblasts embedded in collagen I and topped with a monolayer of mesothelial cells was used to further study cancer cell adhesion and mesothelial displacement. All assays were studied in cells representing different stages of disease progression in the absence or presence of cytostatic doses of mifepristone.

**Results:** 2D in vitro assays demonstrated that the migration and invasive rates of the cells isolated from both patients decreased along disease progression. Conversely, in both patients, cells representing late-stage disease demonstrated a higher adhesion capacity to the 3D organotypic model than those representing an early-stage disease. This adhesive behavior is associated with the in vivo tumor capacity of the cells. Regardless of these differences in adhesive, migratory, and invasive behavior among the experimental protocols used, cytostatic doses of mifepristone were able to inhibit the adhesion, migration, and invasion rates of all cells studied, regardless of their basal capabilities over reductionist or organotypic metastatic in vitro model systems. Finally, we demonstrate that when cells acquire the capacity to grow spontaneously as spheroids, they do attach to a 3D organotypic model system when pre-incubated with conditioned media. Of relevance, mifepristone was able to cause dissociation or “cleavage” of these multicellular structures.

**Conclusion:** Differences in cellular behaviours were observed between reductionist 2D and 3D assays when studying the metastatic capabilities of high-grade serous ovarian cancer cells representing disease progression. Mifepristone inhibited these metastatic capabilities in all assays studied.

## Background

High-grade serous ovarian cancer (HGSOC) is the deadliest histotype of ovarian cancer, accounting for 70-80% of all deaths from the disease [1]. In most cases, HGSOC is diagnosed at late stages, when the disease has already metastasized to distant organs within the peritoneal cavity. HGSOC most commonly metastasizes by transcoelomic dissemination during which cells detach from the primary tumor, either as single cells or as multicellular aggregates, and use the natural flow of peritoneal fluid to reach target tissues [2]. Organs in the peritoneal cavity are covered by a protective layer of peritoneal tissue, made up of a submesothelial stroma composed of fibroblasts and extracellular matrix (ECM), and topped with a monolayer of mesothelial cells. Mesothelial cells form tight junctions between one another and serve a protective function [3]. Two-dimensional reductionist in vitro assays are the standard methods to study the metastatic abilities of cancer cells in vitro (i.e.. adhesion, migration, and invasion) [4]. However, communication between mesothelial cells, fibroblasts, and HGSOC plays an important role in the adhesion and invasion of cancer cells to the peritoneal tissue. To address this process, there has been a rise in the development of organotypic models of disease composed of a minimum of two different cell lines and ECM, with the goal of mimicking microenvironments and physical components that more closely resemble those observed in patients [5]. For the study of ovarian cancer, organotypic models mimicking both the peritoneal tissue and omentum are mostly used, as these two sites are known to often be affected by late-stage disease [2]. Peritoneal tissue models typically contain fibroblasts embedded in collagen I, as ovarian cancer cells have the highest level of adhesion and invasion in the presence of this ECM component [6]; such fibroblasts are topped with a monolayer of mesothelial cells. Tri-dimensional organotypic models of disease have shown, in many cancers, to provide different results than 2D reductionist assays in terms of cellular behavior or in drug responses [7–11].

Treatment options for ovarian cancer have remained stagnant for the last four decades, since the establishment and acceptance of platinum-taxane combination chemotherapy in the 1980s. Although the current standard of treatment, debulking surgery followed by platinum-taxane combination chemotherapy, has a 70-80% initial response rate, most patients will relapse with a chemoresistant disease. The 5-year survival rate after relapse is less than 48% [1]. For this reason, developing new treatment options for these patients is of upmost importance. Previous work in our laboratory has demonstrated the potential of mifepristone (MF), a synthetic steroid that acts both, as anti-glucocorticoid and anti-progestin, as a potential treatment option for ovarian cancer. We demonstrated that MF blocks the growth of ovarian cancer cells [12] and prevents their repopulation after chemotherapy [13] by blocking cells in the G1 phase of the cell cycle as a consequence of increasing the levels of Cdk inhibitors and thus reducing the activity of Cdk2 [14]. We have also shown that MF has antimetastatic potential and can slow the adhesion of highly metastatic cancers, including ovarian, breast, prostate and glial, by rearranging the distribution of fibrillar actin [15], while also interfering with their migratory and invasive capabilities [16].

In this work, we demonstrate that differences in cellular behaviours can be observed between reductionist 2D assays and more complex 3D organotypic assays. Two-dimensional adhesion, migration, and invasion assays demonstrated, in two cases of patient-derived HGSOC cell lines, that cells representing an early-stage disease showed higher metastatic potential than those representing late-stage disease. Conversely, an adhesion assay using an organotypic model demonstrated higher adhesion capacities for the cells obtained at more advanced stage, which coincided with their higher tumorigenicity in vivo. Of relevance, MF inhibited the adhesion, migration, and invasion capacities of all cell lines studied regardless of their metastatic capabilities along disease progression.

## Materials and methods

### Cell lines, culture conditions and treatments

The HGSOC cell lines used were established from two patients and represent disease progression: PEO1, PEO4, and PEO6 from a first patient, and PEO14 and PEO23 from a second patient. PEO1 was originally collected from ascites after first treatment with cisplatin, 5-fluorouracil, and chlorambucil. PEO4 was collected from ascites 10 months after the initial treatment. The patient was treated once more with cisplatin and relapsed a final time, at which point PEO6 was collected from ascites. PEO14 was collected from the ascites of a second patient prior to any treatments (i.e. chemonaïve). PEO23 was collected, from the ascites 7 months after initial treatment with cisplatin and chlorambucil [17]. With written consent from Dr. Langdon (Edinburgh Cancer Research Center, Edinburgh, UK), PEO1, PEO4, and PEO6 cell lines were obtained from Dr. Taniguchi (Fred Hutchinson Cancer Center, University of Washington, Seattle, WA, USA). PEO14 and its longitudinally patient-matched pair PEO23 were obtained from Culture Collections, Public Health England (Porton Down, Salisbury, UK). All cells were cultured in RPMI-1640 (Mediatech, Manassas, VA, USA) supplemented with 5% fetal bovine serum (Atlanta Biologicals, Lawrenceville, GA, USA), 5% bovine serum (Life Technologies, Auckland, New Zealand), 1 mM sodium pyruvate (Corning, Corning, NY, USA), 2 mM L-Alanyl-L-Glutamine (Glutagro™, Corning), 10 mM HEPES (Corning), 0.01 mg/ml human insulin (Roche, Indianapolis, IN, USA), 100 IU penicillin (Mediatech), and 100 µg/ml streptomycin (Mediatech). Cell culture was carried out at 37 °C in a humidified incubator with 95% air/5% CO2 in standard adherent plastic plates.

WI38 fibroblasts were obtained from the American Type Culture Collection (ATCC, Manassas, VA, USA) and were cultured in the same medium as the HGSOC cells. LP9 mesothelial cells were obtained from the Coriell Institute for Medical Research (Coriell, Camden, NJ, USA) and were cultured in a 1:1 combination of F-12 containing L-glutamine (Gibco, from Thermo Fisher Scientific, Waltham, MA, USA) and Medium 199 (Corning) and supplemented with 10% FBS and 0.4 μg/ml hydrocortisone (Sigma Chemical Co., St. Louis, MO, USA). Mifepristone (MF; Corcept Therapeutics, Menlo Park, CA, USA) was dissolved in dimethyl sulfoxide (DMSO) at a concentration of 4,655 µM, and stored at -20°C. During treatment, MF was diluted into culture medium to reach a final concentration of 20 µM, which was previously deemed to be cytostatic [18]. The final concentration of DMSO in culture medium was 0.43%; therefore, vehicle treated cells were exposed to 0.43% DMSO diluted in culture medium. Cells were pre-treated with either MF or medium containing DMSO (vehicle) for 72 h prior to plating and their corresponding treatment was maintained throughout the experiment.

### Cell proliferation and doubling times

The doubling times for the five HGSOC cell lines (PEO1, PEO4, PEO6, PEO14 and PEO23) were repeated twice to confirm the accuracy of the results. For each experiment, cells were plated in 6- well plates at a density of 200,000 cells per well and left to attach for 72 hours. This cell density ensured exponential growth of each cell line while preventing the cells from reaching 100% confluence over the course of the experiment. After 72 hours, triplicate cultures were trypsinized, pelleted by centrifugation at 500 *g* for 5 min, and resuspended in the appropriate growth medium. An aliquot of each cell suspension was counted using the Muse™ Cell Analyzer and the Muse™ Count & Viability Assay Kit. A Count & Viability assay was then run on the Muse™ Cell Analyzer and the total number of cells per sample was collected. These results were considered to be the 0 hour time-point. The same procedure was repeated every 24 hours for a total of 96 hours. Doubling times were calculated as previously described [19] using a nonlinear regression analysis on exponentially growing cells.

### Wound Healing Assay

Each HGSOC cell type was plated in 6-well plates at a density of 200,000 cells per well. Once cells reached approximately 80% confluency, a wound was created along the center of the well, using a 1000 µl pipette tip and a ruler as a guide to ensure the wound was straight and reproducible. Multiple images were then immediately taken of each well, along the wound, using an IN480 Series inverted biological microscope (United Scope LLC, Irvine, CA, USA). Cells were incubated at 37 °C for various time points up to 36 hours. At each time point, images of the wounds were once again taken of each well. Using the OMAX Toupview software (United Scope), the wound width was measured four times per image to determine the wound average. Experiments were repeated with a pre-treatment of either 0.4% DMSO or 20 µM of MF for 72 hours.

### Boyden Chamber Assay

#### Migration

Twenty-four hours before the start of the experiment, cell culture medium was replaced with 0.1% FBS-containing medium. The next day, cells were plated in 0.1% FBS-containing medium in the upper chamber of a 6-well Boyden Chamber (BC) plate at a density of 200,000 cells per well. Ten percent serum-containing medium was added to the lower chamber to act as a chemoattractant. Cells were incubated at 37°C for 30 hours, after which non-migratory cells left in the upper chamber were removed using a sterile cotton swab. The upper chamber was washed with phosphate-buffer saline (PBS) and a second sterile cotton swab was used to ensure that all cells had been removed. Migrated cells were fixed with 4% PFA solution. Twenty 20x field images were taken of each insert by fluorescent microscopy using a Leica DMi8 inverted fluorescence microscope and a Leica LAS X software (Leica Microsystems Canada, Concord, ON). Cells were counted in each image and the average number of cells per 20x field was calculated. Experiments were repeated with a pre-treatment of either 0.4% DMSO or 20 µM of MF for 72 hours.

#### Invasion

To study the invasion of HGSOC cells, the BC was performed in the same way as described for migration. However, 24 hours before cells were plated, inserts were coated with a layer of extracellular matrix (ECM) (Sigma). The stock solution of ECM (9.11 mg/ml) was thawed at 4°C and diluted to a working concentration of 0.6 g/ml. Wells were coated with 500 µl of diluted ECM gel and incubated at 37°C overnight. The next morning, excess ECM gel was removed and the rest of the experiment was performed in the same manner as the migration assay.

### Visualization of migrating and invading cells using cytochemical double fluorescence staining

To improve the visualization of cell migration, cells were stained with a combination of Alexa Fluor®-594 Phalloidin (Life Technologies, Carlsbad, CA, USA) and SYTOX® Green (Molecular Probes, Eugene, OR, USA) or DAPI (Life Technologies), to stain the cytoskeleton and the nucleus respectively. Cells were permeabilized with 0.1% Triton X-100 in PBS for 5 min. To reduce background staining, cells were then incubated in a PBS solution containing 1% BSA for 20 min. The stock solution of Alexa Fluor® 594 Phalloidin was diluted from its 6.6 µM solution in PBS containing 1% BSA at a 1:40 ratio (5 µl stock solution in 200 µl PBS). Cells were incubated with diluted Alexa Fluor® 594 Phalloidin for 20 min. During the last 10 minutes of incubation, cells were exposed to either 4 µl of 50 µM SYTOX® Green (5 mM stock solution) or 300 nM (5 mg/ml stock solution) of DAPI (Life Technologies, Carlsbad, CA) in PBS containing 1% BSA. Finally, the cells were washed with PBS and stored at 4°C.

### Adhesion Assay to Fibronectin

Twenty-four hours before the start of the experiment, 12-well plates were coated with a layer of fibronectin (Gibco). A stock solution of fibronectin (1.0 mg/ml) (Sigma) was thawed at 4°C and diluted to a working concentration of 6 µg/ml in PBS. The next morning, excess fibronectin was removed from the wells and all wells were blocked with a PBS solution containing 1% BSA. Cells were then plated at a density of 100,000 cells/well and left to incubate for 0.5, 1, or 2 hours. The cells were washed twice with PBS, fixed with 100% methanol for 30 min, and stained with a filtered solution of 0.5% (w/v) crystal violet (Sigma) for 10 min. Ten 20x images were taken using an IN480 Series inverted biological microscope (AmScope) and the average number of adherent cells per 20x field was calculated for each well. Experiments were repeated with a pre-treatment of either 0.4% DMSO or 20 µM of MF for 72 hours.

### Organotypic Culture Model

#### Adhesion Assay

The peritoneal tissue is the main site for EOC metastatic lesions; therefore, an organotypic model was utilized based on the model developed by Kenny et al. [6]. To study the adhesive capacity of the five HGSOC cell lines in a tri-dimensional (3D) environment, an organotypic model composed of WI38 fibroblasts embedded in collagen I topped with a monolayer of LP9 mesothelial cells was created. Four thousand WI38 cells were suspended in a solution of 2 ml of LP9 culture medium and 30 µg of collagen I (Corning), per single-well chamber slide. The fibroblasts were incubated in a humidified environment at 37°C and 5% CO2 for 4 hours. Three-hundred and fifty thousand LP9 mesothelial cells were plated per chamber slide and incubated in a humidified environment at 37°C and 5% CO2 overnight, to form a confluent monolayer. HGSOC were incubated for 45 min with 5 µM of CellTracker™ Deep Red Dye (Invitrogen) diluted in serum-free medium. Stained HGSOC cells were then plated at a density of 250,000 per chamber slide and left to adhere to the LP9 monolayer for 2, 4, or 24 hours. Cells were fixed with 4% PFA and stored at 4°C. Experiments were repeated with a pre-treatment of either 0.4% DMSO or 20 µM of MF for 72 hours.

#### Conditioned Medium

Organotypic culture models were created as previously described and incubated with conditioned medium from all 5 HGSOC cell lines for 24 hours. Medium was collected from each cell culture after incubating with the cancer cells for 24 hours. Organotypic models were fixed with 4% PFA and stored at 4°C.

### Immunofluorescence

Cells were permeabilized using 0.1% Triton X-100 for 5 min. To reduce background staining, the cells were then incubated with a PBS solution containing 1% BSA for 20 min. Cells were incubated with the primary antibody diluted in 0.2% BSA at 4°C overnight (see Additional File 1: Table S1 for the source and specific concentrations of each antibody). Cells were washed twice with PBS for 3 min and then incubated with the secondary antibody for 30 min. Cells were washed twice with PBS for 3 min and stored at 4°C. Images were taken using a Leica DMi8 inverted fluorescence microscope and a Leica LAS X software (Leica Microsystems) or a Cytation™ 3 Cell Imaging Multi-Mode Reader with Gen5 software (Biotek, Winooski, VT, USA).

### Statistical Analysis

All data represent means ± s.e.m. and statistical significance was defined as p<0.05. One-way analysis of variance (ANOVA) followed by Bonferroni’s test was used to compare the means among three different cell lines. Two-way ANOVA followed by Bonferroni’s test was used to compare the means of groups receiving different treatments over time. Unpaired student’s *t*-test was used when comparing the means between two different cell lines.

## Results

### High-grade serous ovarian cancer cells representing early-stage disease have a shorter doubling time than those representing late-stage disease

Proliferation assays were performed for each cell line. The doubling times of cell lines representing an early-stage disease, PEO1 and PEO14, were found to be shorter than those of cell lines representing late-stage disease, PEO4, PEO6, and PEO23. This difference was most obvious between PEO1, PEO4, and PEO6, with PEO6 growing almost three times slower than PEO1. The difference between PEO14 and PEO23 was less drastic, but still showed an increased doubling time related to disease progression (Fig. 1A, B). The morphology of each cell line in culture also varies along disease progression. PEO1 and PEO14, cells representing an early-stage disease, tend to grow linearly without any overlap. As the disease progresses, cells begin to grow more in a 3D fashion, forming foci as cells begin to overlap one over the other. This is particularly obvious in PEO6 and PEO23 where an evident elevation in the cell monolayer can be visualized. PEO4 demonstrates a phenotype between PEO1 and PEO6 with the beginning of a 3D growth, however not as evident as in PEO6 or PEO23.

**Fig. 1.**
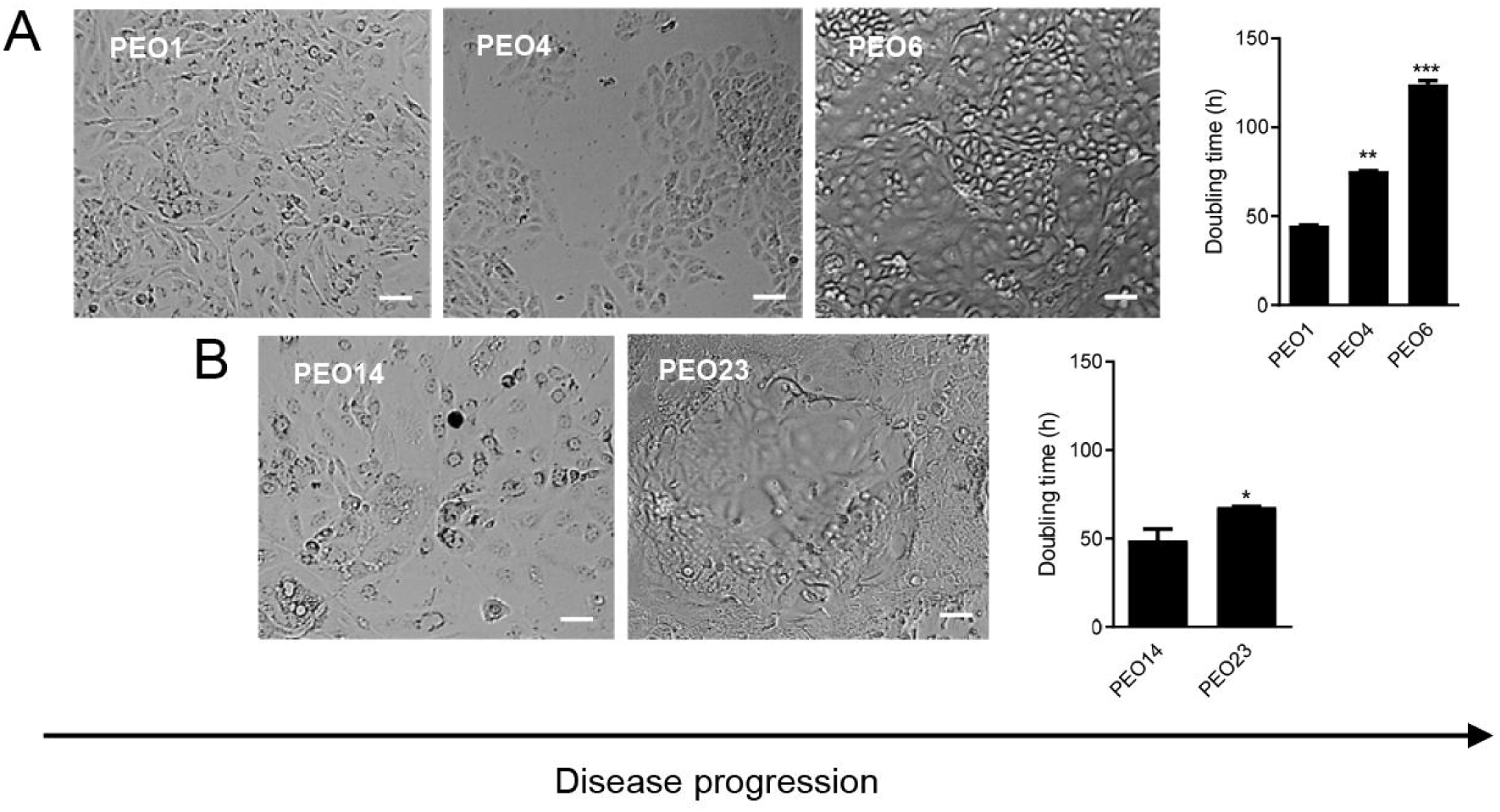
Doubling-times of HGSOC cells obtained along disease progression. The number of cells was collected every 24 hours for a total of 96 hours. Doubling times were calculated using a nonlinear regression analysis on exponentially growing cells. For panels **(A)** and **(B)**, data shown represents the mean ± s.e.m. Panels **(C) – (G)** are brightfield images representing the morphologies of each HGSOC cell in culture. Scale bars = 50 μm.

### The migratory capacity of high-grade serous ovarian cancer cells decreases along disease progression as assessed by wound healing and Boyden chamber assays

The migration capacity of each cell line was studied via two separate methods, the wound healing and Boyden chamber assays, which we previously optimized using fluorescence microscopy [16]. To ensure that in the wound healing assay we were assessing cellular migration and not cellular proliferation, all time points studied were less than the doubling time of each cell line. In addition, we confirmed that indeed if proliferation occurred, it was mostly away from the wound, not at the wound site as assessed by staining with phospho-histone H3, a marker of mitosis (Additional File 2: Fig. S1). When compared to each other, PEO1 cells were found to have a higher migration rate than the PEO4 and PEO6 counterparts isolated later in the disease. After 36 hours, PEO1 closed 40% of the wound width, while PEO4 and PEO6 migrated to close 20% and 10% of the wound respectively (Fig. 2 A, [i-iv]). This decrease in migration rate along disease progression was also found to be true between the PEO14 and PEO23 pair. PEO14 was able to close 35% of the wound while PEO23 closed approximately 20% of the wound during the same time (Fig.2 B [i-iii]).

**Fig. 2.**
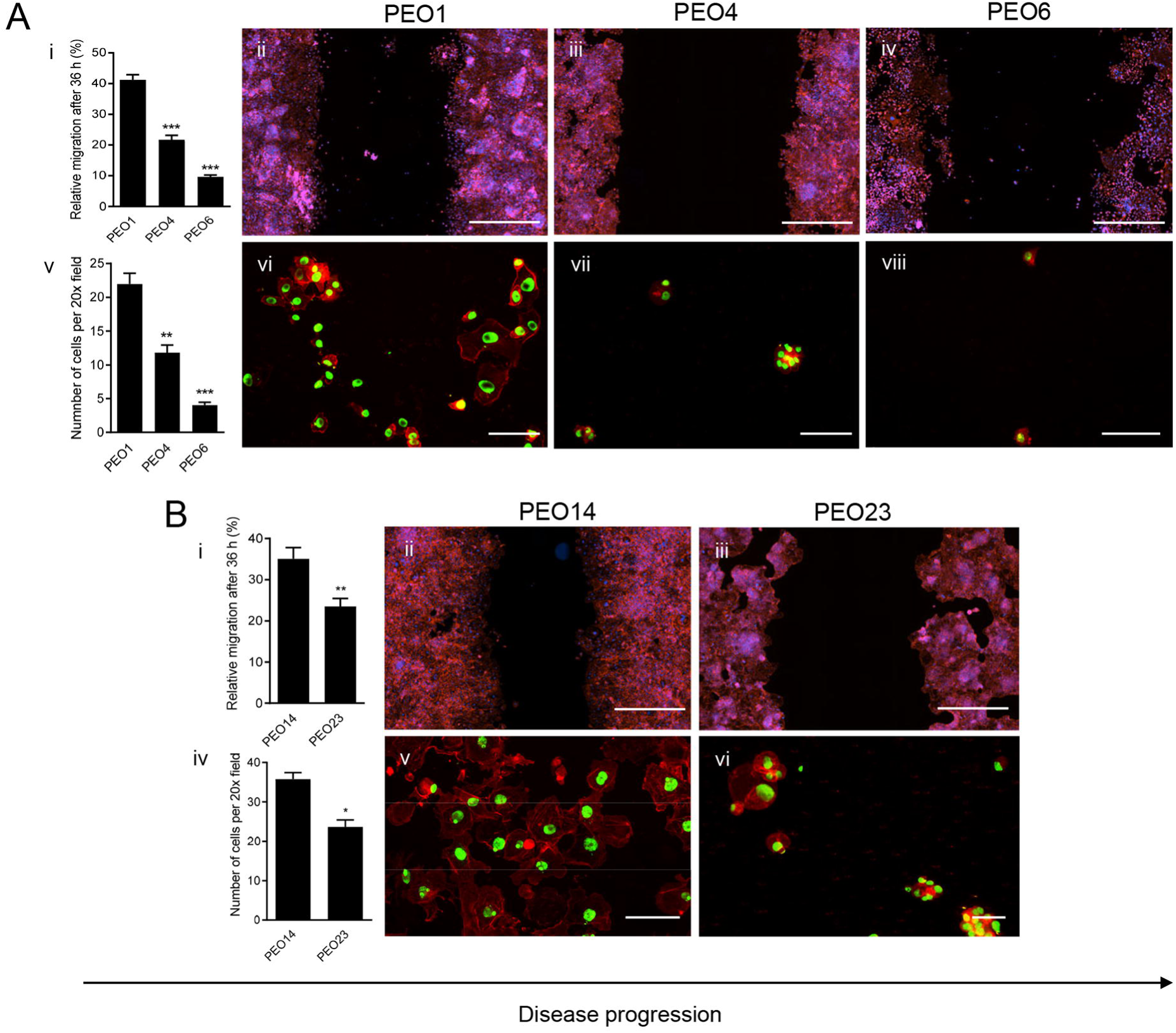
Migration rates of HGSOC cells along disease evolution in both wound healing (**A [i-iv], B [i-iii]**) and Boyden chamber (**A [v-viii], B [iv-vi]**) assays. In the wound healing assay, multiple images were taken per wound at 0 and 36 hours and the wound width was measured 4 times per image. The difference between the initial and final wound widths was calculated and converted into percentage of wound closure. Panels **(A [ii-iv])** and **(B [ii-iii])** are visual representations of each cell line after 36 hours, labelled with DAPI, to stain the nucleus, and Alexa Fluor-594 Phalloidin, to stain the cytoskeleton. Scale bar = 1000 μm. In the Boyden chamber assay, cells that had migrated through the insert after 30 hours were counted. Panels (**A [vi-viii]**) and (**B [v-vi]**) are visual representations of migrated cells after 30 hours, labelled with SYTOX green, to stain the nucleus, and Alexa Fluor-594 Phalloidin, to stain the cytoskeleton. Scale bar = 100 μm. Data shown represents the mean ± s.e.m. For panels (**A[i]**) and (**A[v]**), **P<0.01, ***P<0.001 compared to PEO1, and ###P<0.001 compared to PEO6. Statistical analysis was done using one-way ANOVA followed by Bonferroni’s test. For panels (**B[i]**) and (**B[iv]**), *P<0.05, **P<0.01 compared to PEO14. Statistical analysis was done using paired student *t-*test.

Boyden chamber assays were performed to confirm this decrease in migration capacity along disease evolution. Once again, when the series of cell lines were compared to each other, it was found that cell lines representing an early-stage disease had a higher migration rate than cell lines representing a late-stage disease. After 36 hours, PEO1 cells were able to cross the membrane, occupying a large portion of the surface, while PEO4 and PEO6 were found much more scarcely after having been provided the same amount of time to migrate (Fig, 2 A [v-viii]). Similar results were obtained between the second series of cell lines. PEO14 was found to be the most migratory out of all the cell lines, occupying almost the entirety of the membrane surface after 36 hours. PEO23 migrated less rapidly than PEO14 and covered less of the surface of the membrane (Fig. 2 B [iv-vi]).

### Cells representing early-stage disease undergo simple cell migration whereas cells representing late-stage disease undergo collective cell migration

Cancer cell migration has long been described as individual cells undergoing physical changes to move from a primary to a distant site. However, it has recently been shown that two distinct migration patterns exist; single cell migration, during which cells migrate to surrounding tissues independently of one another, and collective cell migration, during which cells cluster together and migrate as a collective unit to distant sites [20]. Through the addition of a double fluorescence labeling of the actin cytoskeleton and nucleus, variations in the migration pattern between cells representing early-stage disease and cells representing late-stage disease were uncovered. PEO1 cells seem to be undergoing single-cell migration, as demonstrated by cells separating from one another into the wound, in the wound healing assay, and covering most of the membrane surface, once migrated, in the Boyden chamber assay (Fig.3A [i, iii]). PEO4 cells were observed closing the wound, in a wound healing assay, as a sheet of cells, and were found to form tight clusters when migrating in the Boyden chamber assay, suggesting a collective cell migration pattern (Fig. 3A [ii-iv]). The observation that cells representing early-stage disease undergo single-cell migration while cells representing a late-stage disease undergo collective cell migration was further supported in the second series of cell lines. PEO14 cells seem to separate from one another as they progress into the opening of the wound, in a wound healing assay, and cells having migrated through the pores in a Boyden chamber assay spread throughout the well, often individually from one another (Fig. 3B [i, iii]). PEO23 cells demonstrated a migration pattern similar to that of PEO4 cells, moving as a cohesive sheet of cells in the wound healing assay and forming distinct clusters of cells after migrating through the pores in the Boyden chamber assay (Fig. 3B [ii-iv]). The studies with PEO6 cells were not considered as there were not enough migrating cells in the Boyden chamber assay to reach a conclusion (data not shown).

**Fig. 3.**
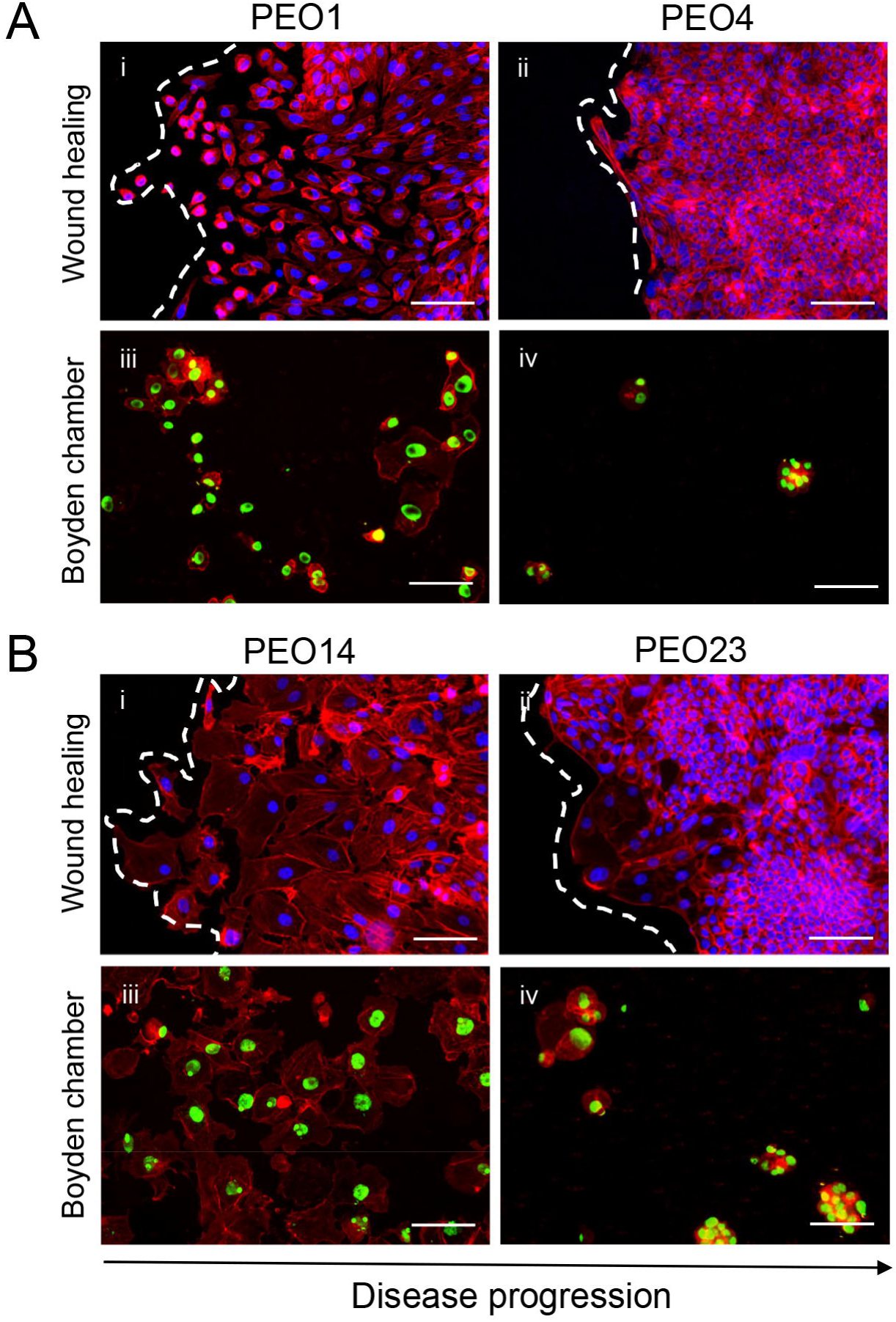
Staining wound healing and Boyden chamber assays with a double cytochemical labeling allowed for the observation of varying migration patterns along disease progression. Cells were labelled with Alexa Fluor-594 Phalloidin, to stain for the cytoskeleton, and either DAPI (**A[i-ii], B[i-ii]**) or SYTOX green (**A[iii-iv], B[iii-iv]**), to stain for the nucleus. Scale bar = 100 μm. White dashed lines represent the front of the wound.

### The invasive capacity of high-grade serous cell lines decreases along disease progression

The invasive capacity of each cell line was determined through the Boyden chamber assay, similar to the migration assay; however, plates were incubated with a layer of ECM overnight, challenging the cells to secrete metalloproteases and chew through the ECM before migrating through the membrane pores. There was found to be a decrease in invasive capacity along disease progression. PEO1 was found to be twice as invasive as PEO4 after 30 hours of invasion, with PEO6 being slightly less invasive than PEO4 (Fig. 4A). These results were confirmed in the second case of cell lines. Once again, PEO14 was the most aggressive cell line, invading twice as rapidly as PEO1 after 30 hours, while the invasive capacity of PEO23 was half that of PEO14 (Fig. 4B).

**Fig. 4.**
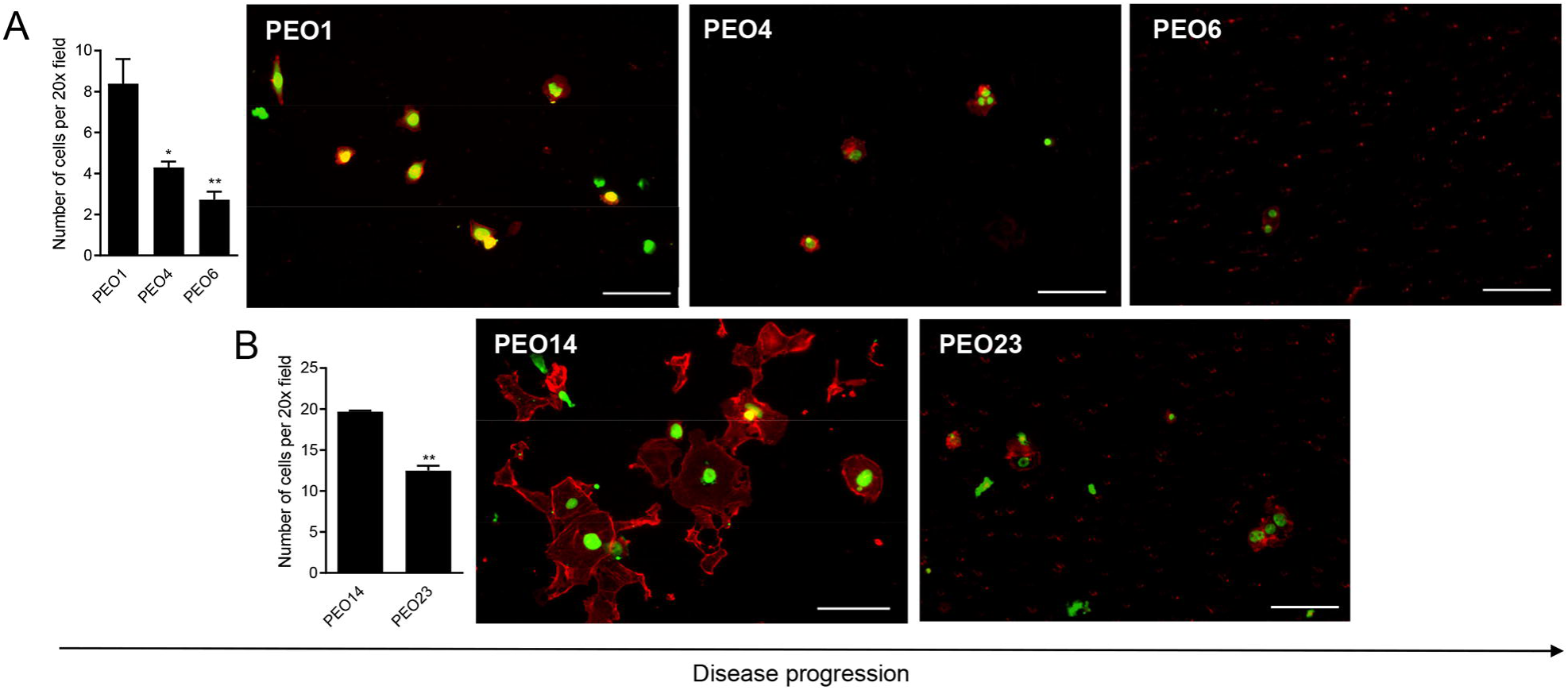
The invasion rate of HGSOC cells across disease progression in a Boyden chamber assay. Boyden chamber assays were performed similarly to the migration assays but with the addition of a layer of extracellular matrix. Panels (**A[ii-iv]**) and (**B[ii-iii]**) are visual representations of invaded cells after 30 hours, labelled with SYTOX green, to label the nucleus, and Alexa Fluor-594 Phalloidin to label the cytoskeleton. Scale bar = 100 μm. Data shown represents the mean ± s.e.m. For panel (**A[i]**), *P<0.05, **P<0.01 compared to PEO1. Statistical analysis was done using one- way ANOVA followed by Bonferroni’s test. For panel (**B[i]**) **P<0.01 compared to PEO14. Statistical analysis was done using paired student *t*-test.

### The expression of cell adhesion molecules E-cadherin, N-cadherin and CD44 varies along disease progression while cells migrate in a wound healing assay

The ability of cancer cells to migrate and invade has long been associated with the ability of cells to undergo epithelial-mesenchymal transition (EMT), in which cells go from a structural phenotype associated with high levels of E-cadherin, to a more migratory and invasive one, in which N-cadherin is upregulated [21]. The expression of E-cadherin and N-cadherin varies not only between HGSOC cell lines, but also within each cell line itself. For instance, PEO1 demonstrates few clusters of E-cadherin positive cells surrounded by a majority of individually arranged N-cadherin positive cells at the wound (Fig. 5A [i]). However, PEO1 cells away from the wound shows a heterogeneous pattern with various E-cadherin clusters surrounded by N-cadherin positive cells (Fig. 5A [iv]). PEO4, a cell line found to be less migratory than PEO1, demonstrated a much more epithelial phenotype, with most cells being E-cadherin positive, both at (Fig. 5A [ii]) and away (Fig. 5A [v]) from the wound; furthermore, several PEO4 cells were found to be positive for both E-cadherin and N-cadherin. PEO6 cells at the wound were found to be represented by a mix of positive N-cadherin cells, negative for both antigens, and only few E-cadherin positive cells (Fig. 5A [iii]). Away from the wound (Fig 5A [vi]) cells were mainly isolated, with low E-cadherin expression, with few positive clusters (Fig.5A [vi]). For the second set of paired cell lines (Fig. 5B), although PEO14 was found to be the most migratory cell line in both the wound healing and Boyden chamber assays, it demonstrated a highly epithelial phenotype; many cells are double positive for E-cadherin and N-cadherin, while others are only E-cadherin positive at (Fig. 5B [i]) and away (Fig. 5B [iii]) from the wound. PEO23, its later stage counterpart, was found to be entirely E-cadherin positive both at (Fig 5B [ii]) and away (Fig.5B [iv]) from the wound.

**Fig. 5.**
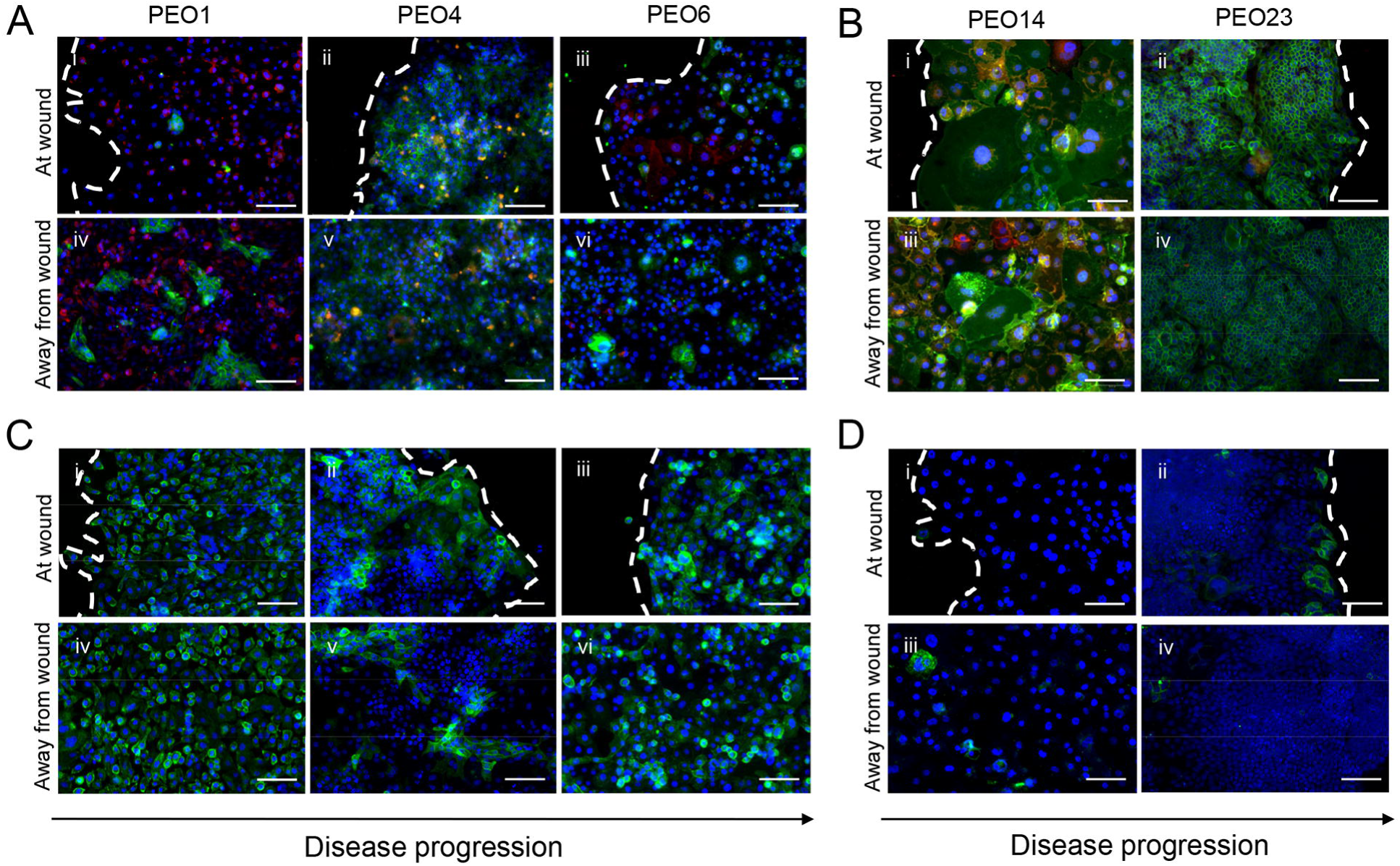
Immunofluorescence staining for E-Cadherin, N-Cadherin, and CD44 shows heterogeneity along disease progression and within the same cell line. Wound healing assays were performed as previously described. After 36 hours, cells were fixed with 4% PFA and stained for E-cadherin (green), N-cadherin (red), and DAPI (blue) in panels (**A**) and (**B**) and for CD44 (green) and DAPI (blue) in panels (**C**) and (**D**). Scale bar = 100 μm. White dashed lines represent the front of the wound.

CD44 is an adhesion molecule shown to bind to the ECM component, hyaluronic acid (HA). The binding of CD44 to HA has been shown to promote ovarian cancer cell binding to peritoneal tissue and ovarian cancer cell migration [2]. PEO1 was found to be entirely positive for CD44, both at the wound (Fig. 5C [i]) and away from the wound (Fig. 5C [iv]). PEO4 started to demonstrate more heterogeneity in CD44 positivity, with patches of positive cells surrounded by negative cells. Interestingly, clusters of cells migrating into the wound were found to be CD44 positive (Fig. 5C [ii]) when compared with the patches observed away of the wound (Fig. 5C [v]). The CD44 pattern of PEO6 veered back to resembling that of PEO1, with most of the cells being positive for the marker, with a few negative cells in both at the wound (Fig. 5C [iii]) or away from the wound (Fig. 5C [vi]). Surprisingly, PEO14 was found to be almost completely negative for CD44, both at the wound (Fig. 5D [i]) and away from the wound (Fig. 5D [iii]). PEO23 demonstrated a distinct pattern of clustered CD44 positive cells right on the edge of the wound (Fig. 5D [ii]). Conversely, cells away of the wound were negative for CD44 (Fig. 5D [iv]). In summary, we show that expression of E-cadherin, N-cadherin, and CD44 are highly variable among cells representing disease progression, many times even showing differences between cells found at the wound when compared with cells located away from the wound in this particular migration assay.

The ability of cancer cells to adhere to a distant site is an important process in the end phase of metastasis. Of interest, however, when we assessed the capacity of the cells to adhere to plates pre-coated with fibronectin, we found that in the case of the PEO1/4/6/ or PEO14/23 cellular series despite they adhere within 2 hours to the plates, there is no much difference in the adhesion capacity among the cell lines; only PEO14 resulted to have superior adhesive capacity to fibronectin when compared to all other cell lines studied (Additional File 3: Figure S2).

### The adhesive capacity of high-grade serous ovarian cancer cells increases along disease evolution in an organotypic model system composed of collagen-embedded fibroblasts topped with mesothelial cells

The peritoneal wall is composed of a submesothelial stroma topped with a monolayer of mesothelial cells and is the main target for HGSOC metastasis. To study further the metastatic capacity of HGSOC cells along disease progression, an organotypic model composed of fibroblasts embedded in collagen I and topped with a monolayer of mesothelial cells was developed. Contrary to what we observed in previous section using 2D migration and invasion assays, we found, based on the adhesion capacity to this organotypic model system that the cells seem to have more aggressive metastatic behavior with disease progression. Adhesion of HGSOC cells to the organotypic model demonstrated an increase along disease progression in both PEO1/4/6 and PEO14/23 cellular series. In the first series, PEO6 cells were found to adhere most efficiently to the monolayer of mesothelial cells when compared to early-stage counterparts, PEO1 and PEO4 (Fig. 6A [i]). Immunofluorescence staining of the mesothelial monolayer allowed for the visualization of changes in the mesothelial cells over time. After 24 hours of incubation, the mesothelial cell layer remains mainly intact after the adhesion of PEO1 cells; however, in the presence of PEO4 and PEO6 cells the mesothelial monolayer was displaced over time, and ovarian cells were found to adhere within the spaces left by the LP9 cells (Fig. 6A [iii, iv, vi, and vii]). In the second cellular series, we observed a similar pattern in adhesive capacity, with PEO23 being more adhesive to the organotypic model system than PEO14 (Fig. 6B [i]). Opposite to the first cellular series, in the second series, PEO14, representing an early-stage disease, managed to displace the monolayer of LP9 cells, while PEO23 simply adhered to the surface of the mesothelial cell monolayer, leaving it intact (Fig. 6B [iii-v]).

**Fig. 6.**
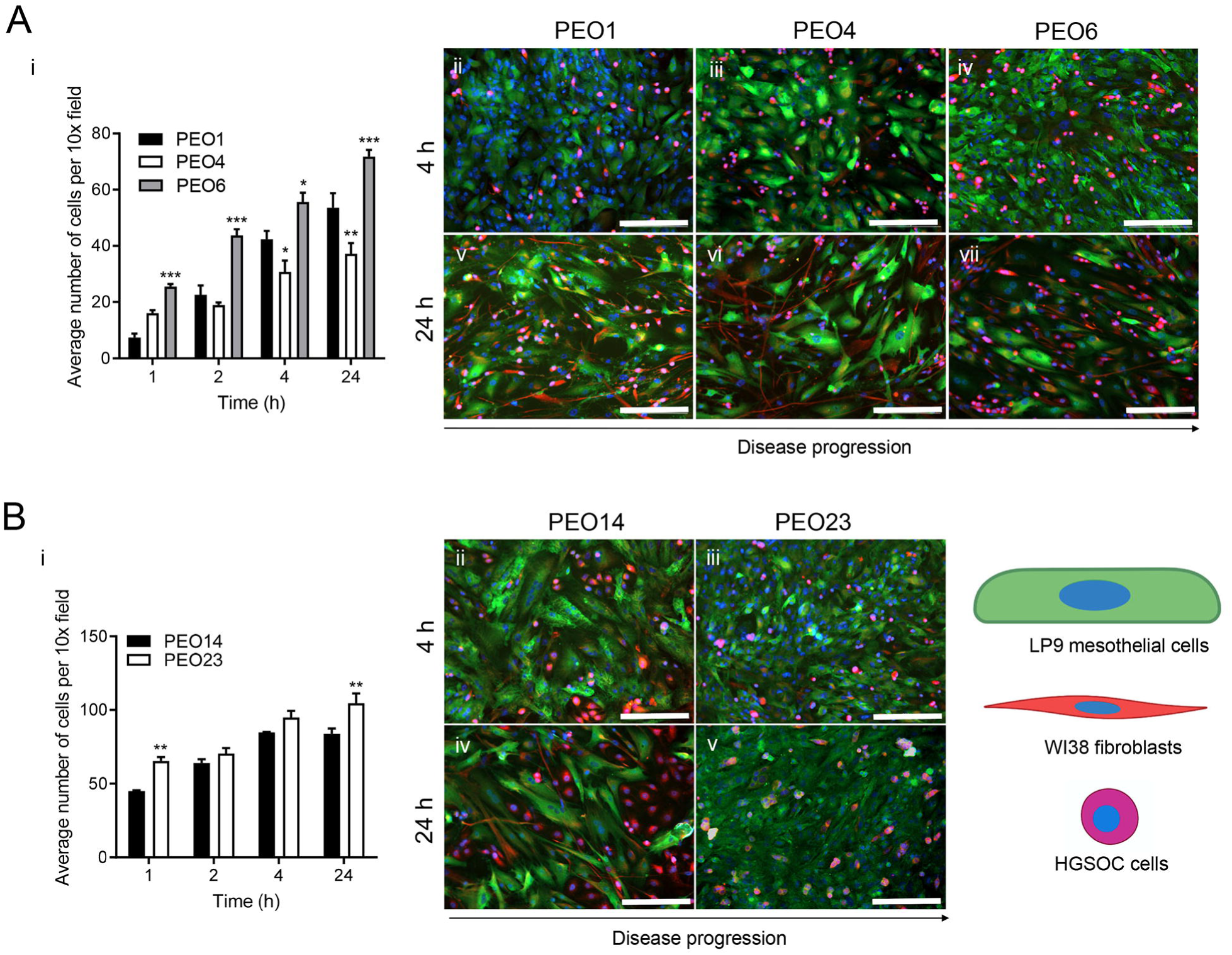
The adhesion rate of HGSOC cells to organotypic models, composed of fibroblasts embedded in collagen I topped with a monolayer of LP9 cells, increases along disease progression. Panels (**A[ii-vii]**) and (**B[ii-v]**) are a visual representation of HGSOC cells adhered to the LP9 monolayer after 4 hours (**A[ii-iv], B[ii-iii]**) and 24 hours (**A[v-vii], B[vi-v]**). HGSOC cells were incubated with Cell Tracker™ Deep Red before plating. Cells were fixed with 4% PFA and stained for calretinin (green, [mesothelial cells]) and vimentin (red, [fibroblasts]) by immunofluorescence, and DAPI, to stain the nucleus. Scale bar = 200 μm. Data shown represents the mean ± s.e.m. For panel **(A[i]),** *P<0.05, **P<0.01, ***P<0.001 compared to PEO1. For panel **(B[i]),** **P<0.01 compared to PEO14. Statistical analysis was done using two-way ANOVA followed by Bonferroni’s test.

In the previous figure, we observed that LP9 cells, when incubated with many of HGSOC cells, seem to be displaced. To determine if the displacement of LP9 mesothelial cells is due to the HGSOC cells or to secreted factors, we incubated the organotypic model system containing only fibroblasts and mesothelial cells with the conditioned media of the HGSOC cells for 24 hours. Conditioned media from PEO4, PEO6, and PEO14 cells caused major displacement of LP9 cells, whereas conditioned media from PEO1 and PEO23 caused minor displacement of LP9 cells (Additional File 4: Fig. S3). To further confirm that a secreted factor is responsible for the displacement of LP9 cell monolayer, PEO23 cells that did not cause major displacement were incubated in conditioned media from PEO14 cells that did cause large LP9 displacement. Results shown in Additional File 5: Fig. S4 clearly indicate that the PEO14 conditioned media caused displacement of LP9 cells while at the same time significantly decreased the adhesion of PEO23 cells.

### HGSOC spheroids adhere to an organotypic model only in the presence of conditioned media

The formation of spheroids is an important step in the survival and dissemination of HGSOC cells. HGSOC cells are commonly found as spheroids within patient ascites, and it is thought the cell-cell interaction favors their survival within the peritoneal fluid. To assess the metastatic capacity of HGSOC spheroids adhering to an organotypic multicellular model, PEO6 cells were used. PEO6 is a cell line with the capacity to form floating spheroids spontaneously (Fig. 7A), a phenomenon that we amply described previously [22]. The ability for PEO6 spontaneously formed spheroids to adhere to the organotypic model was assessed for 24 hours. Spontaneously-formed PEO6 floating spheroids resuspended in fresh media and incubated for 24 hours, were not able to adhere (Fig.7B, inset), and were not able to cause mesothelial cell displacement; only a few individual cells were observed adhering to the mesothelial monolayer (Fig. 7B). However, when resuspended and incubated in PEO6 conditioned media (i.e. the media from a confluent culture of PEO6 cells), the PEO6 spheroids were able to adhere to the LP9 monolayer while causing their displacement (Fig 7C).

**Fig. 7.**
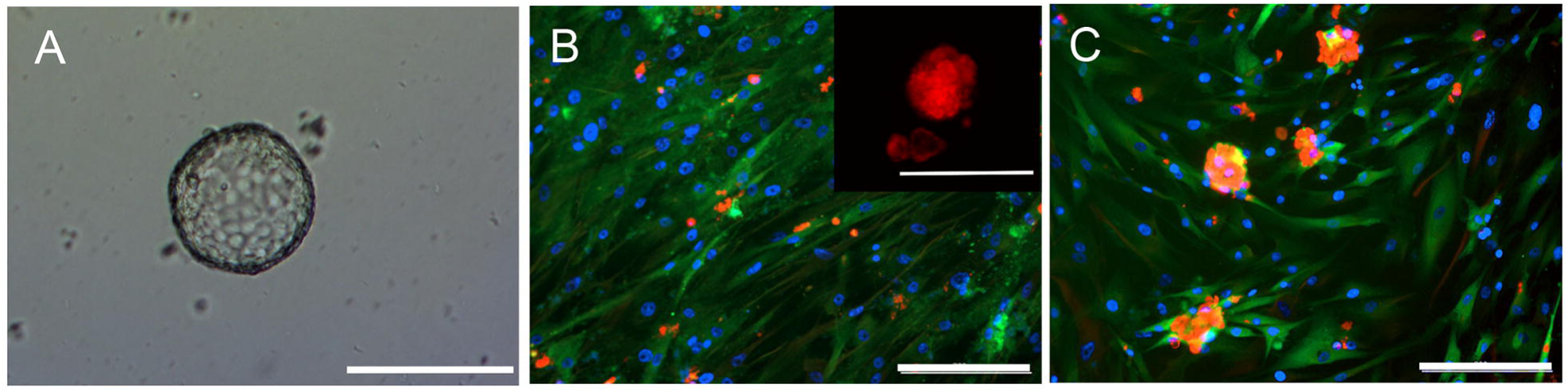
HGSOC spheroids do not adhere to an organotypic model composed of fibroblasts embedded in collagen I topped with a monolayer of LP9 mesothelial cells, nor cause mesothelial cell displacement unless incubated with conditioned media. Panel **A** represents a brightfield image representing the morphology of PEO6 spheroids. Panel **B** is a visual representation of PEO6 spheroids adhered to the organotypic model after 24 hours. Panel **(C)** represents PEO6 spheroids incubated with adherent PEO6 conditioned media for 24 hours. Cells were fixed with 4% PFA and stained for calretinin (green, [mesothelial cells]), vimentin (red, [fibroblasts]) by immunofluorescence, and DAPI (blue), to stain the nuclei. Inset in panel **B** represents spheroids that are not attached (floating) when incubated in non-conditioned media. Scale bar = 200 µm.

### Mifepristone inhibits adhesion, migration, and invasion of high-grade serous ovarian cancer cells regardless of their stage in the disease

Adhesion, migration, and invasion are three key processes important for cancer cells to perform in order to metastasize from a primary to a distant site. Therefore, therapeutic options targeting these processes are essential. HGSOC cells were treated with a previously determined cytostatic dose of MF for 72 hours [12]. When challenged to adhere to fibronectin coated plates, MF-treated HGSOC cells were significantly slower to adhere in both cases of cell lines, regardless of their stage in the disease. Even PEO14, which had the highest capacity to adhere to fibronectin, was inhibited by MF to the same extent as the other cell lines (Fig. 8A [i-iii], and Fig. 8B [i-ii]). When MF-treated cells were challenged to migrate through a Boyden chamber assay, once again the drug was able to inhibit the migration rate of each cell line, regardless of their basal differences in migratory capacity (Fig. 8A [iv-vi], and Fig. 8B [iii-iv]). Wound healing assays were also repeated with MF-treated cells and demonstrated similar results (Additional File 6: Fig. S5). Finally, in the Boyden chamber assays repeated with the addition of a layer of ECM, once again MF was able to inhibit the invasion of both lines of HGSOC cells, regardless of their basal invasive abilities (Fig. 8A [vii-ix], and Fig. 8B [v-vi]).

**Fig. 8.**
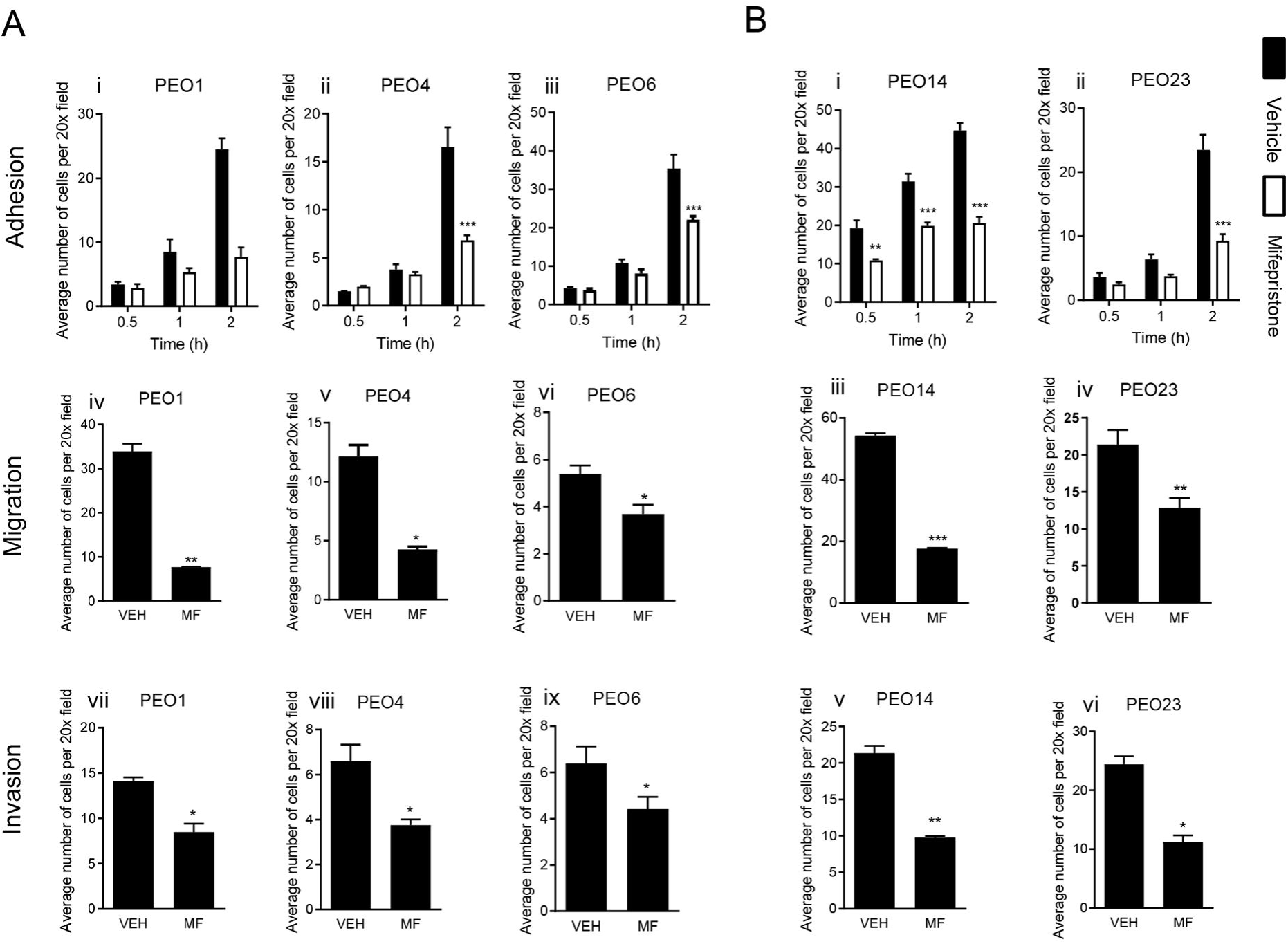
Mifepristone inhibits the adhesion, migration, and invasion capacity of all five HGSOC cell lines despite basal capabilities. HGSOC cells were treated with 20 μM of MF for 72 hours prior to plating. Panels (**A[i-iii]**) and (**B[i-ii]**) represent adhesion assays to fibronectin. Panels (**A[iv-vi]**) and (**B[iii-iv]**) represent Boyden chamber migration assays. Panels (**A[vii-ix]**) and **(B[v-vi]**) represent Boyden chamber invasion assays. Data shown represents the mean ± s.e.m. For panels (**A[i-iii]**) and (**B[i-ii]**), **P<0.01, and ***P<0.001compared to Vehicle. Vehicle (closed bars), MF (open bars). Statistical analysis was done using two-way ANOVA followed by Bonferroni’s test. For panels (**A[iv-ix]**) and (**B[iii-vi]**), *P<0.05, **P<0.01 compared to Vehicle. Statistical analysis was done using student *t*-test.

### Mifepristone inhibits the adhesion of high-grade serous ovarian cancer cells in an organotypic model system

HGSOC cells pretreated with cytostatic doses of MF were challenged to adhere to an organotypic model system composed of fibroblasts embedded in collagen I, topped with a monolayer of mesothelial cells. Once again, regardless of basal capacities of each cell line, MF was able to slow the rate of adhesion of each cell line to the organotypic model (Fig. 9 A [i-iii], and Fig. 9 B [i-ii]).

**Fig 9.**
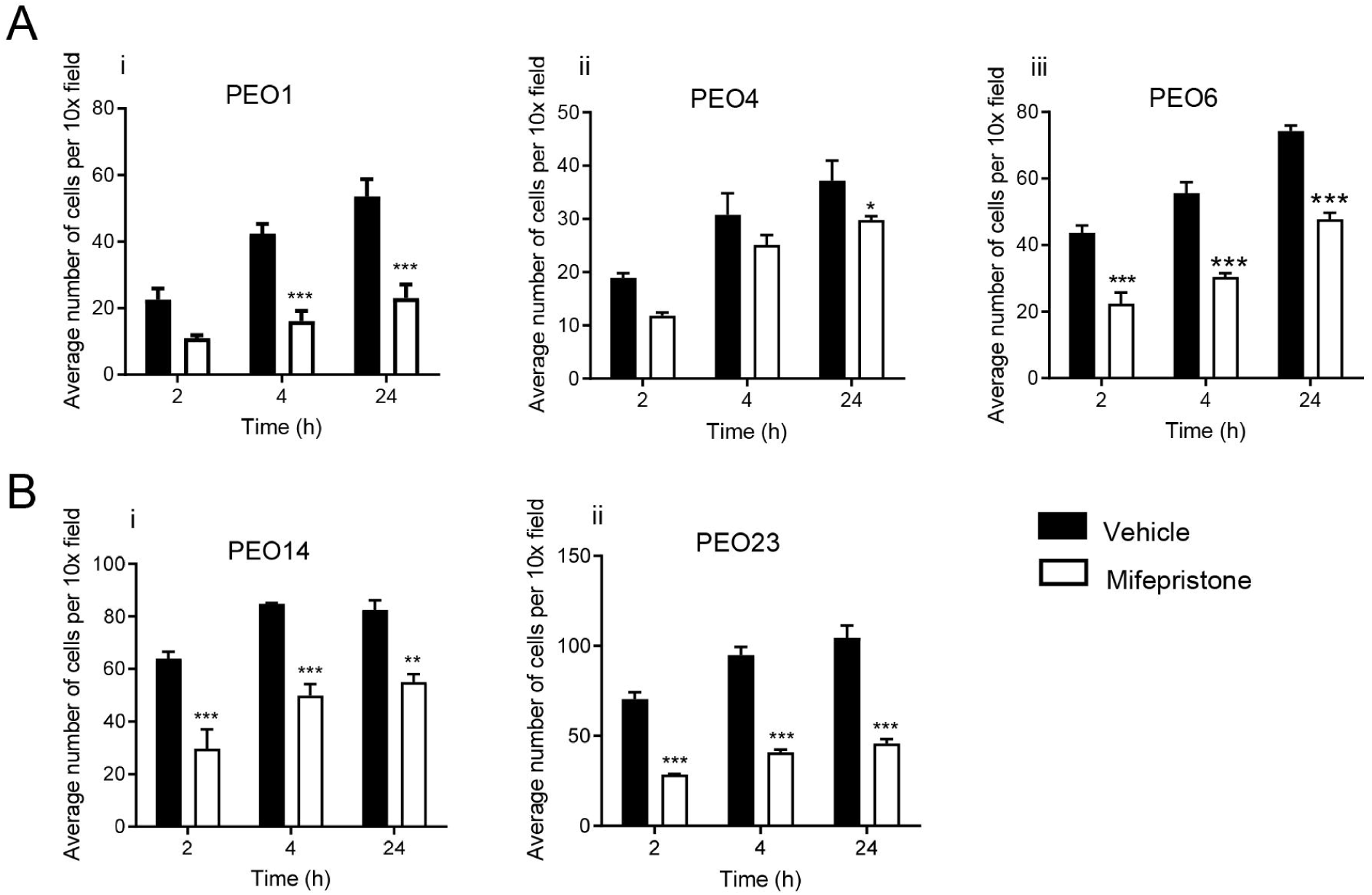
Mifepristone inhibits the adhesion of all five HGSOC cell lines to an organotypic model, composed of fibroblasts embedded in collagen I topped with a monolayer of LP9 cells. HGSOC cells were treated with 20 μM of MF for 72 hours prior to plating. Data shown represents the mean ± s.e.m. For panels (**A[i-iii]**) and (**B[i-ii]**), **P<0.01 and ***P<0.001 compared to Vehicle. Vehicle (closed bars), MF (open bars). Statistical analysis was done using two-way ANOVA followed by Bonferroni’s test.

### Mifepristone promotes the dissociation of spheroids adhered to the mesothelial cells

We previously showed that spontaneously formed PEO6 spheroids were able to adhere to a monolayer of mesothelial cells when in the presence of conditioned media (Fig. 7). When we cultured the spheroids in the presence of conditioned media containing MF, the spheroids are smaller and have a tendency to dissociate into individual cells as demonstrated by the reduced quantity of un-dissociated spheroids and the increased number of isolated cells (Fig. 10 A, B). Spheroids and small clusters of dissociated cells seem to adhere over the mesothelial cells but not over the empty spaces created by PEO6-conditioned media in the absence or presence of MF.

**Fig. 10.**
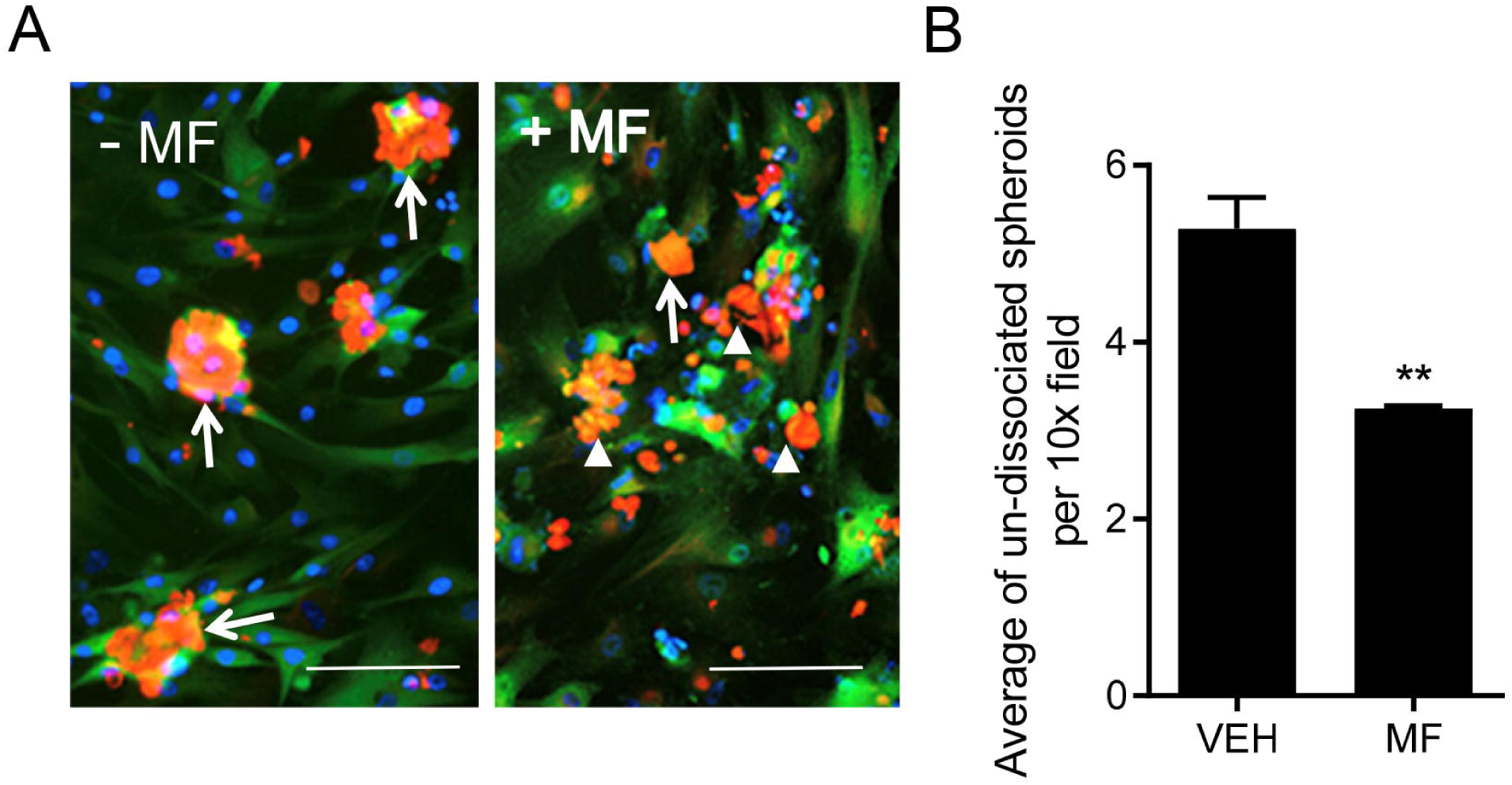
Mifepristone promotes the dissociation of PEO6 spheroids, incubated in adherent PEO6 conditioned media, to the organotypic model composed of fibroblasts embedded in collagen I and topped with a layer of mesothelial cells. PEO6 spheroids were treated with 20 μM of MF for 72 hours prior to plating. Scale bar = 200 µm. Data shown represents the mean ± s.e.m. For panel **C**, **P<0.01 compared to Vehicle. Statistical analysis was done using student *t*-test.

## Discussion

In this work, we characterized the metastatic abilities of two series of HGSOC cell lines, representing disease progression, and studied the ability of cytostatic doses of MF to inhibit these capabilities. The majority of ovarian cancer cases are diagnosed once the disease has already metastasized, emphasizing the importance of understanding and treating this phenomenon. Metastasis can be divided into three complex processes: adhesion, migration, and invasion. Migration involves the rearrangement of the cytoskeleton leading to an elongated morphology and increased contractility [4]. Two types of cellular migration have been described: single cell and collective cell migration. In single cell migration, each individual cell undergoes physical changes before migrating. Single cell migration is often associated with EMT, in which E-cadherin is often downregulated indicating the loss of an epithelial phenotype [20]. Collective cell migration describes the ability of cells to migrate as clusters, in which cells retain their cell-cell junctions and migrate as a collective group. This allows for migrating cells to carry along cells with a more epithelial phenotype, all while maintaining contact with one another. Collective cell migration has been associated with a more invasive phenotype in many cancers including breast [23], thyroid [24] and prostate [25], with leading cells having a more mesenchymal phenotype than those in the rear. Contrary to what we expected, our results demonstrated that cells representing a late-stage disease were less migratory than their early-stage counterparts in 2D in vitro assays. However, it was also noted that PEO4 and PEO23 (both representing advanced disease) were found to be more E-cadherin positive than PEO1 and PEO14 (representing earlier stage disease), demonstrating a more epithelial phenotype as the disease progressed. Furthermore, PEO4 and PEO23 obtained at advanced disease were observed migrating as clusters of cells, suggesting that they undergo collective cell migration. These observations combined suggest that HGSOC may not undergo what has long been believed to be the classic form of metastasis, in which invasive cells undergo EMT, developing a more mesenchymal phenotype to enhance migration and invasion, before adhering and invading a distant site [26]. Instead, although cells undergoing single cell invasion migrate more rapidly, cells undergoing collective cell migration are more efficient, as the crosstalk between cells helps to coordinate the direction of the migration of the group [27]. This could explain the more rapid migration rates found in cells representing early-stage disease (i.e. PEO1 and PEO14) and the observation that these seem to be undergoing single cell migration, although they may be less efficient to metastasize into a 3D organotypic model system. Due to collective cell migration being associated with increased metastatic potential, the ability of HGSOC cells to travel also as spheroids in the peritoneum could contribute to the aggressivity and low survival rate of the disease compared to other cancers.

The upregulation and downregulation of various adhesion molecules is important throughout the entire process of cancer metastasis and is involved in triggering conformational changes within cancer cells as well as determining the ECM protein cells preferentially adhere to [28]. HGSOC has been shown to have a predisposition for the peritoneal tissue, composed of a single layer of mesothelial cells atop of sub-mesothelium of ECM, composed of many proteins including collagen I and fibronectin [29]. Although both ECM proteins play a role in ovarian cancer metastasis, studies have shown that ovarian cancer cells adhere more readily to collagen I than fibronectin [5], as well as demonstrate an increased ability to migrate when cultured on collagen I and a decreased ability when cultured on fibronectin [30]. This coincides with the differences observed between simple adhesion assays on fibronectin and organotypic models containing collagen I. Adhesion to fibronectin showed minimal differences in adhesion rates between cell lines (except for the only chemonaïve cell line, PEO14). It was only when plated onto the more complex organotypic models that differences between the adhesive capabilities of cell lines began to be observed. In this case, there was a clear increase in adhesion along disease progression in both cell line groups. It has been shown that single cell migration is associated with weak adhesion to ECM, while collective cell migration involves strong cell-cell junctions and interactions with underlying ECM in order for the cellular group to move forward [31]. This could explain the reason for the differences observed in adhesion to the organotypic model, as cells with the higher adhesion rates seems to be cells with a collective cell migration pattern.

Hyaluronic acid (HA) is another major ECM protein that binds the adhesion molecule CD44 and has been implicated in ovarian cancer metastasis; increased levels have been associated with poor ovarian cancer outcome [32, 33]. Immunofluorescent staining of CD44 in a wound healing assay showed that both PEO4 and PEO23, cells representing a late-stage disease, showed CD44 positive cells when migrating into the wound, presumably to facilitate adhesion. This pattern is consistent with a collective cell migration pattern, in which leading cells are primed to undergo migration and are the first to adhere. PEO1 and PEO14, cells representing an early-stage disease, demonstrated extremely different levels of CD44. All PEO1 cells were found to be CD44 positive, while the majority of PEO14 cells were found to be CD44 negative, which is somewhat controversial as both represent early-stage disease with the only difference that PEO1 are chemosensitive whereas PEO14 are chemonaïve.

In this study, we demonstrated that behaviors of HGSOC cells differ between 2D simplistic in vitro adhesion assays vs. 3D organotypic in vitro adhesion assays. Organotypic models have the goal of providing cells with an artificial tumor microenvironment, which is not present in standard in vitro assays. Cell-cell communication between mesothelial cells, fibroblasts, and ovarian cancer cells are relevant to achieve metastasis to the peritoneal tissue. For example, cancer-associated fibroblasts promote HGSOC metastasis [34, 35], while mesothelial cells inhibit the adhesion and invasion of HGSOC cells [36]. Therefore, the presence of both cell types is important in observing HGSOC cell behaviors more closely resembling those encountered in patients. Adhesion assays in organotypic models showed an increase in adhesion rate along disease progression in both series of cell lines. Furthermore, through the addition of an immunofluorescence staining with calretinin to label the mesothelial and vimentin to label the fibroblasts, a displacement of mesothelial cells was observed. This is concordant with what it was found by Kenny et al. that showed that mesothelial cells are absent under the peritoneal tumor mass in wide spread ovarian cancer disease [37]. In our results, PEO4, PEO6, and PEO14 were all observed disrupting the mesothelial cell monolayer after 24 hours of incubation. Although mesothelial cell displacement was not dependent on disease progression in this case, it was associated with tumorigenicity in vivo. Our laboratory recently demonstrated that only PEO4, PEO6, and PEO14 are capable of forming tumors in vivo, while PEO1 and PEO23 show no signs of tumorigenicity even after 14 months of inoculation into the peritoneal cavity of immunosuppressed mice [22]. The disruption of the mesothelial monolayer is an important step of HGSOC metastasis, as cancer cells can adhere more easily to the underlying ECM than to the mesothelial cells themselves. The exact mechanism for mesothelial cell displacement is still unknown, however it was suggested that it involves the removal of mesothelial cells by ovarian cancer cells using force [38]. Of interest, when organotypic models were incubated with conditioned media from PEO4, PEO6, or PEO14 cells for 24 hours but without the cancer cells, the LP9 monolayer was found to be disrupted in a similar fashion as when the cells were present and adhered. These results suggest the presence of secreted factors that could be communicating with the mesothelial cells, promoting their displacement. It has been shown that the presence of pro-inflammatory cytokines can cause structural changes to mesothelial cells. Particularly, tumour necrosis factor-α (TNF-α), interleukin-1 ß (IL-1ß), and hepatocyte growth factor (HGF) have all been shown to cause mesothelial cell retraction, exposure of the underlying ECM, while facilitating the adhesion of cancer cells to the mesothelial cell monolayer [39–42]. Transforming growth factor-ß1 (TGF-ß1) has also been suggested to be involved in the progression of peritoneal metastasis by causing fibrosis of mesothelial cells, decreasing the integrity of the mesothelium, and increasing the secretion of chemokines, homing HGSOC cells to the peritoneal tissue [43]. Furthermore, exposure to various pro-inflammatory cytokines causes variations in the level of adhesion molecules present on the surface of mesothelial cells. Interestingly, IL-1ß has been shown to increase levels of CD44 while TNF-α causes an opposite effect [39].

The formation of HGSOC spheroids is thought to serve a protective function while traveling in the peritoneal fluid, increasing cell survival and resistance to anoikis [44, 45]. As the spheroids reach the peritoneal tissue, communication between mesothelial cells and HGSOC cells primes the environment, promoting adhesion and invasion of cancer cells [46, 47]. PEO6 spheroids demonstrated increased adhesion capacity when exposed to adherent PEO6 conditioned media, implying the requirement of secreted factors for the spheroids to adhere to the organotypic system representing a reductionist model mimicking the peritoneal wall.

The effect of MF on the adhesion, migration, and invasion abilities of each cell line was studied in this work. It was found that regardless of the variations in metastatic abilities of each cell type, MF was able to inhibit the migration and invasion in wound healing and Boyden chamber assays, adhesion to fibronectin, and to the 3D organotypic model system. The ability of MF to inhibit these properties was not dependent on disease progression nor platinum-sensitivity, making it a good potential treatment candidate for progressive HGSOC disease. The exact mechanism behind this inhibition is still unknown; however, a previous report from our laboratory demonstrated that MF inhibits the adhesion of cancer cells representing ovarian, breast, prostate, and nervous system cancers, through the redistribution of fibrillar actin into cortical actin ruffles that are not adherent, thus diminishing the surface of cells with adhesion capacity [15]. This is a mechanism that could also affect the ability of cells to migrate and invade, as these processes involve front to rear polarization and actin cytoskeletal rearrangement, in order for cells to move forward [27]. When lamellipodia cannot establish themselves firmly, they tend to retract towards the center of the cell, halting the migration process [48]. Stress fibers are essential for the adhesion of cells to the substratum, as well as the morphological changes undergone during migration. The loss of stress fibers in MF-treated cells could also be a reason for the reduction in migration capacity [49]. Cells undergoing collective cell adhesion undergo similar actin cytoskeletal rearrangement as cells undergoing single cell migration, just with the addition of adherens junctions between cells [27]. Therefore, this hypothesis for the mechanism of action of MF in the inhibition of migration and invasion would pertain to cells undergoing both single cell and collective cell migration.

Finally, we demonstrate that spheroidal HGSOC structures are dissociated by MF into small clusters and isolated cells in the presence of conditioned media. This suggests that MF may target both individual cells as well as spheroidal structures floating in the peritoneal cavity and prevent their adhesion to the peritoneal wall.

## Conclusions

Two series of HGSOC cell lines representing disease progression were used throughout this study and variations along disease progression were observed. Migration and invasion rates were found to decrease along disease progression, while adhesion to fibronectin was found to be relatively similar between all cell lines except for chemonaïve PEO14 cells. Adhesion to an organotypic model, composed of fibroblasts embedded in collagen I and topped with a monolayer of mesothelial cells, was found to increase along disease progression. The capacity of cell lines to cause clearance of the mesothelial layer was associated with the presence of a secreted factor or factors and with their capacity to create tumors in vivo. Regardless of these differences in behaviors among in vitro approaches to study metastatic behavior, MF was able to inhibit the migration, invasion, and adhesion rates of all cell lines studied, including the capacity to dissociate spheroidal structures into individual cells, which are further prevented from adhesion. These data highlight the complexity of transcoelomic metastasis, and that models used to study metastasis are insufficient to understanding the mechanisms involved. This work highlight that while the mechanism of metastasis of HGSOC cells still need further mechanistic investigation, MF has a strong anti-metastatic effect against this deadly disease regardless of its progression.

## Supporting information

Cells migrating into the wound express no PHH3

Adhesion rate of HGSOC cells to fibronectin

Addition of conditioned media causes displacement of mesothelial cells

Adhesion of PEO23 cells to an organotypic model when incubated with PEO14 conditioned media

Mifepristone inhibits migration of HGSOC cells in a wound healing assay

## Abbreviations

BC: Boyden chamber
BS: bovine serum
ECM: Extracellular matrix
EMT: Epithelial- mesenchymal transition
FBS: Fetal bovine serum
HA: Hyaluronic acid
HGF: Hepatocyte growth factor
HGSOC: High-grade serous ovarian cancer
Interleukin-1ß: IL-1ß
Mifepristone: MF
PHH3: Phospho-histone H3
TGF- ß1: Transforming growth factor-ß1
TNF-α: Tumour necrosis factor-α
VEH: Vehicle
WH: Wound healing.

## Acknowledgements

We thank Drs. Robert Roe and Hazel Hunt (Corcept Therapeutics) for supplying pharmaceutical grade mifepristone.

## Author’s contributions

SJR carried out all experiments. ASMN oversaw and supported the study design. AAG and CMT designed the study, and supervised the graduate student. SJR drafted the manuscript, which is part of her Doctoral Thesis. AAG and CMT contributed to the writing of the final version of the manuscript. All authors read and approved the final manuscript.

## Availability of data and materials

Not applicable.

## Declarations

### Ethics approval and consent to participate

Not applicable

### Consent for publication

Not applicable.

---

**Table S1.**
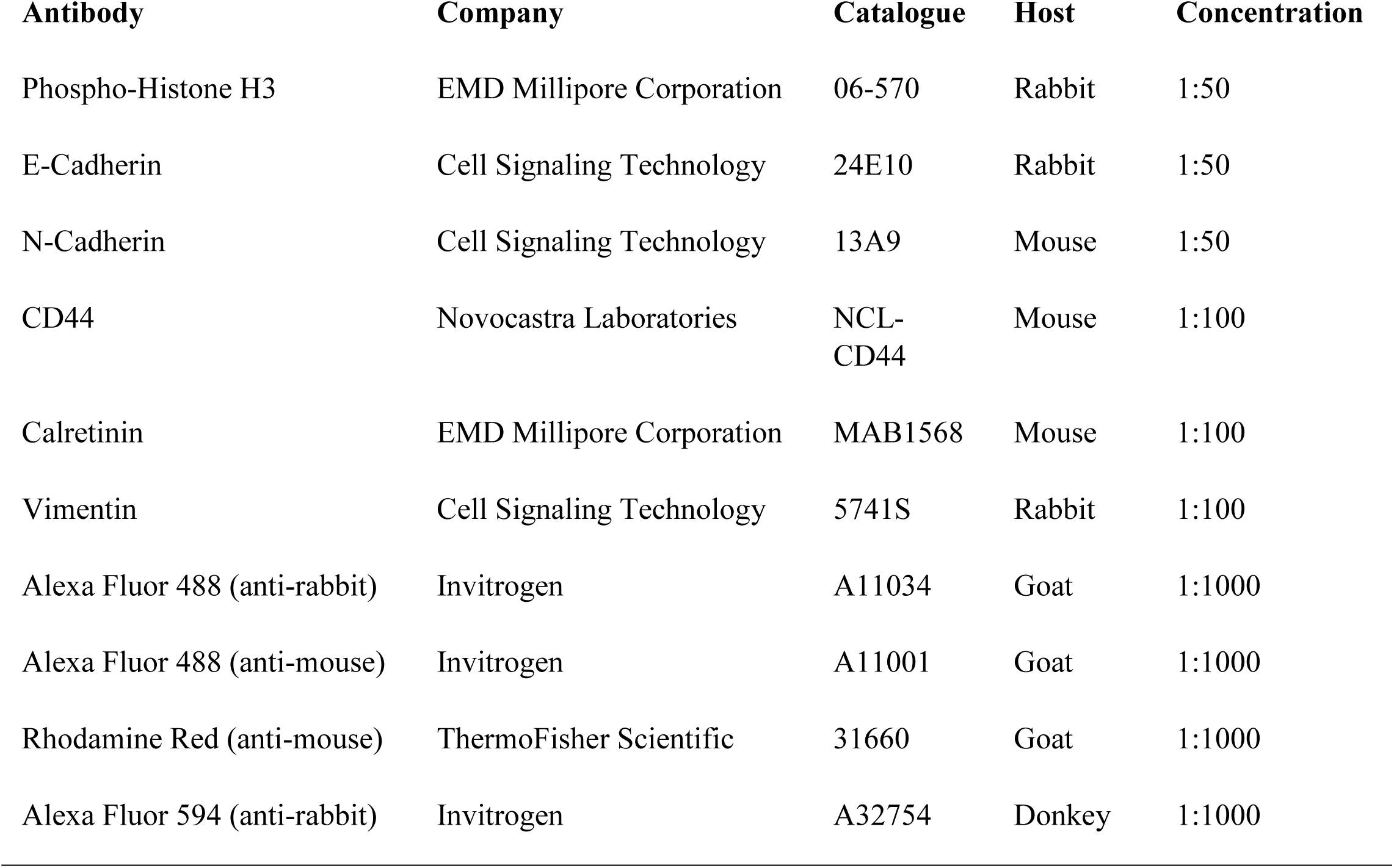
Source and dilutions of antibodies utilized in this work.

Fig. S1 Cells migrating into the wound express no PHH3 compared to an increase in positivity away from the wound, demonstrating that cells are migrating into the wound and not proliferating. Cells were fixed after 36 hours and stained for PHH3 (green) by immunofluorescence, and Alexa Fluor-594 Phalloidin to stain for the cytoskeleton. The white dashed lines represent the front of the wound. Scale bar = 100 μm.

Fig. S2 The adhesion rate of HGSOC cells to fibronectin only changes along disease progression in one case. Plates were pre-coated with fibronectin and cells were left to adhere for 2 hour. Data shown represents the mean ± s.e.m. Statistical analysis was done using one-way ANOVA followed by Bonferroni’s test. ***P<0.001 compared to the other cell lines.

Fig. S3 The addition of conditioned media to organotypic models causes a displacement of the mesothelial cell monolayer in PEO4, PEO6, and PEO14. Panels are a visual representation of organotypic models incubated with conditioned media from the 5 HGSOC cell lines studied, for 24 hours. Cells were fixed with 4% PFA and stained for calretinin (green, [mesothelial cells]), vimentin (red, [fibroblasts]) by immunofluorescence, and DAPI (blue), to stain the nuclei. Scale bar = 200 μm.

Fig. S4 The adhesion rate of PEO23 to an organotypic model, composed of fibroblasts embedded in collagen I topped with a monolayer of LP9 cells, decreases when incubated with PEO14 conditioned media. Panels (**A** and **B**) are a visual representation of PEO23 adhered to the LP9 monolayer incubated without (**A**) or with (**B**) PEO14 conditioned media (C.M.) for 24 hours. PEO23 cells were incubated with Cell Tracker^TM^ Deep Red before plating. Cells were fixed with 4% PFA and stained for calretinin (green, [mesothelial cell]s), vimentin (red, [fibroblast]s) by immunofluorescence, and DAPI (blue), to stain the nuclei. Scale bar = 200 μm. For panel **(C)** **P<0.01 compared to control. Statistical analysis was done using student *t*-test.

Fig. S5 Mifepristone inhibits the migration capacity of all five HGSOC in a wound healing assay, despite basal capabilities. PEO1, PEO4 and PEO6 series, panel (**A[i-iii]**) and PEO14 and PEO23 series, panel (**B [i-ii]**). HGSOC cells were treated with 20 μM of MF for 72 hours prior to plating. Data shown represents the mean ± s.e.m. *P<0.05, **P<0.01, ***P<0.001 compared to Vehicle. Vehicle (closed bars), MF (open bars). Statistical analysis was done using two-way ANOVA followed by Bonferroni’s test.

