## Supplementary figures and images for "The metastatic capacity of high-grade serous ovarian cancer cells changes along disease progression: inhibition by mifepristone"

### Addition of conditioned media causes displacement of mesothelial cells

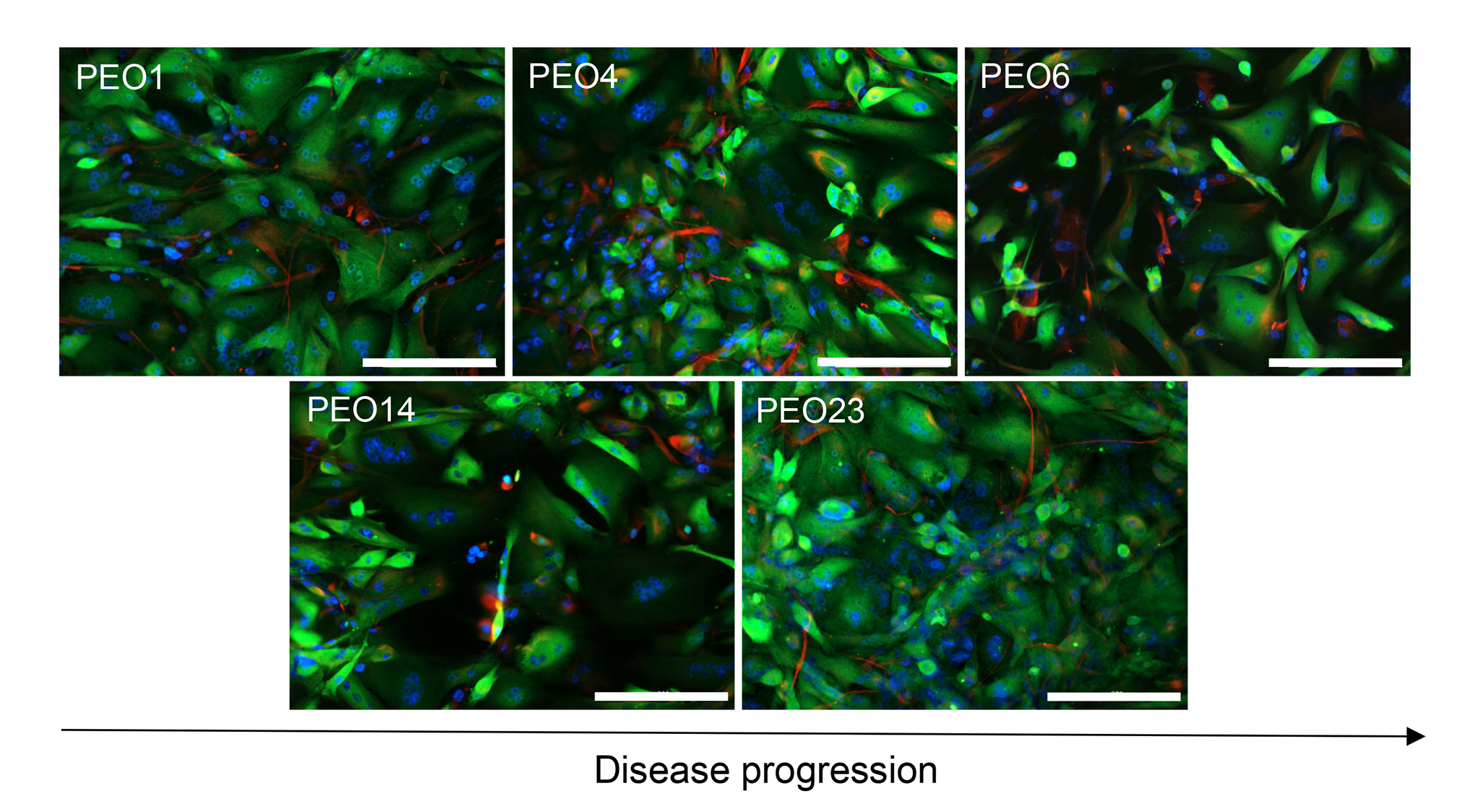

### Adhesion of PEO23 cells to an organotypic model when incubated with PEO14 conditioned media

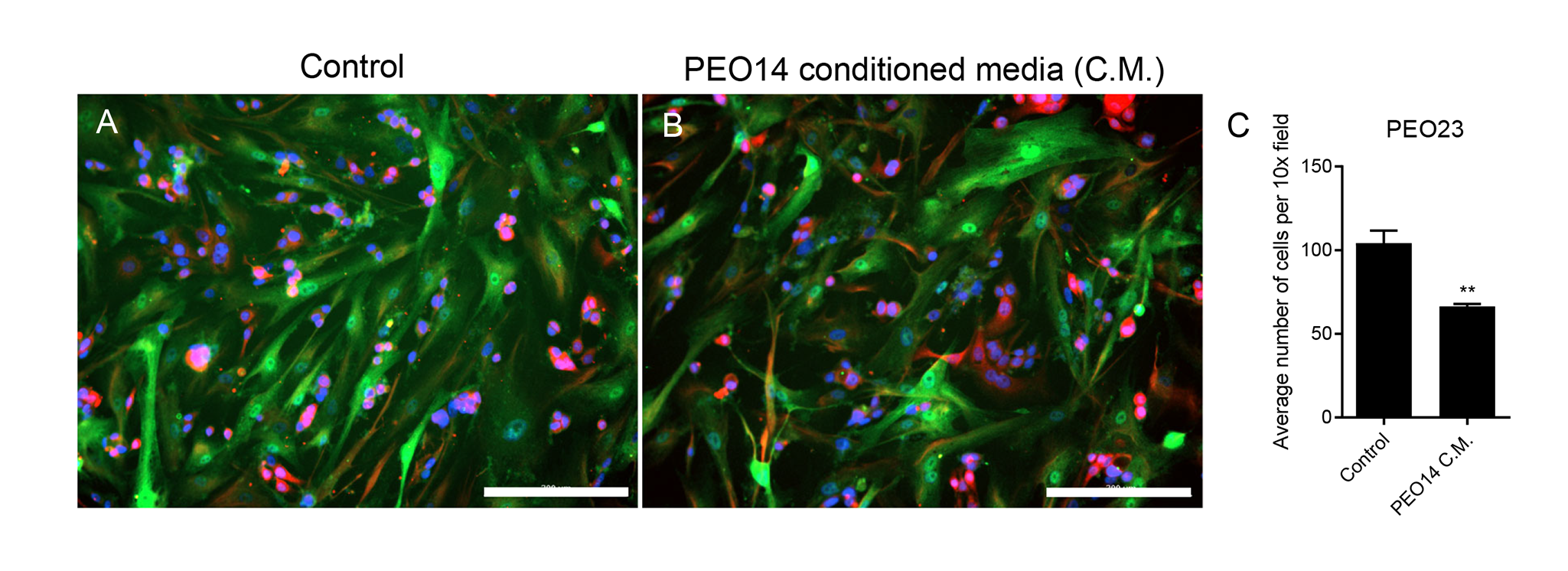

### Adhesion rate of HGSOC cells to fibronectin

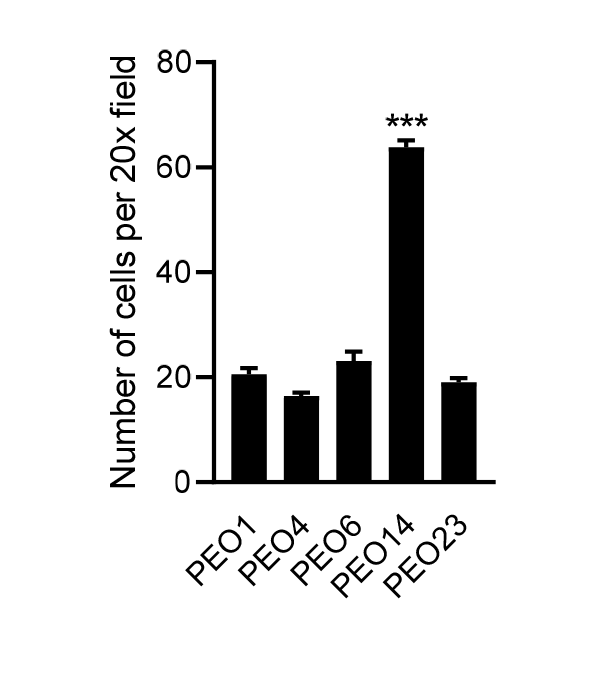

### Cells migrating into the wound express no PHH3

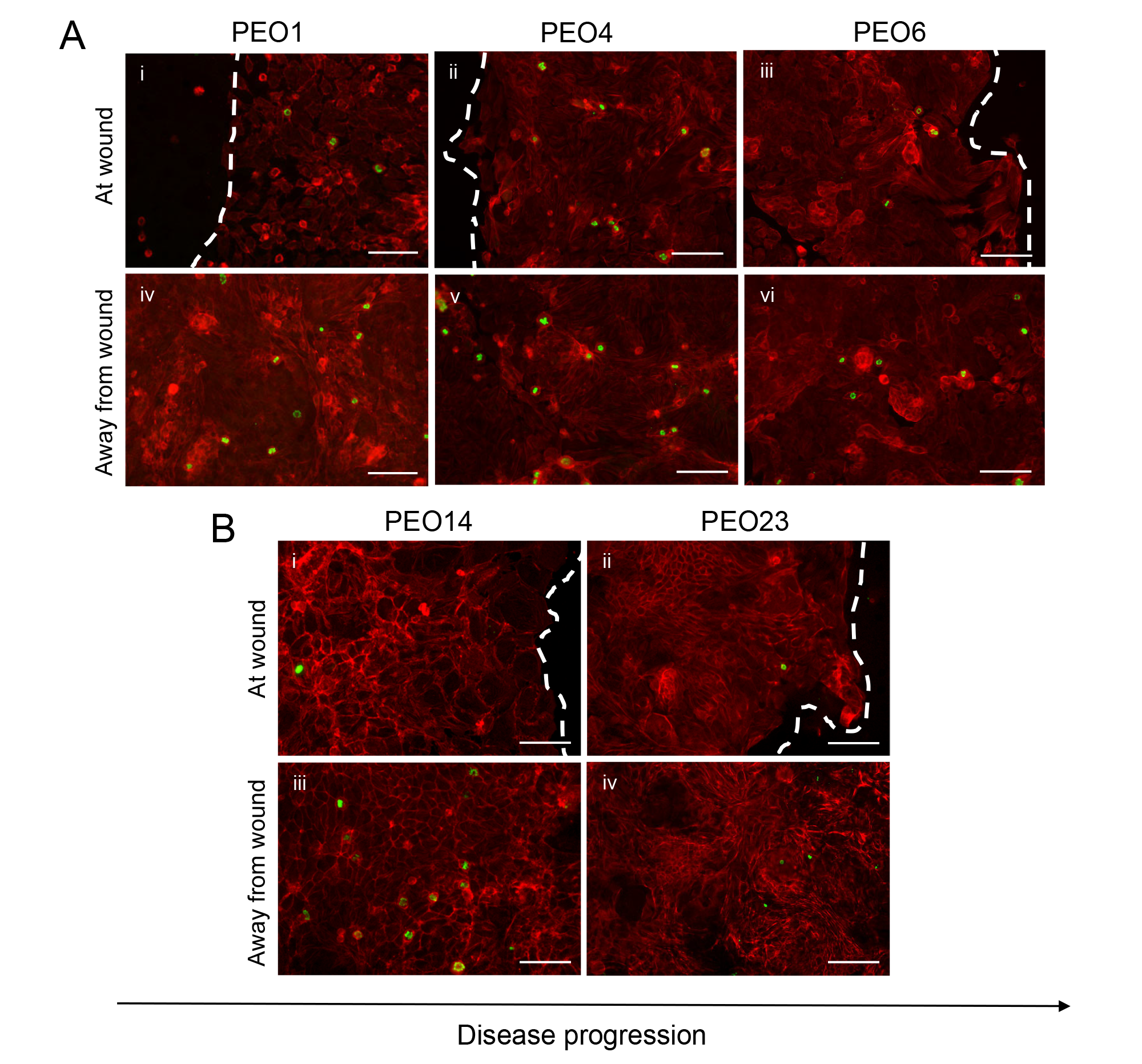

### Mifepristone inhibits migration of HGSOC cells in a wound healing assay

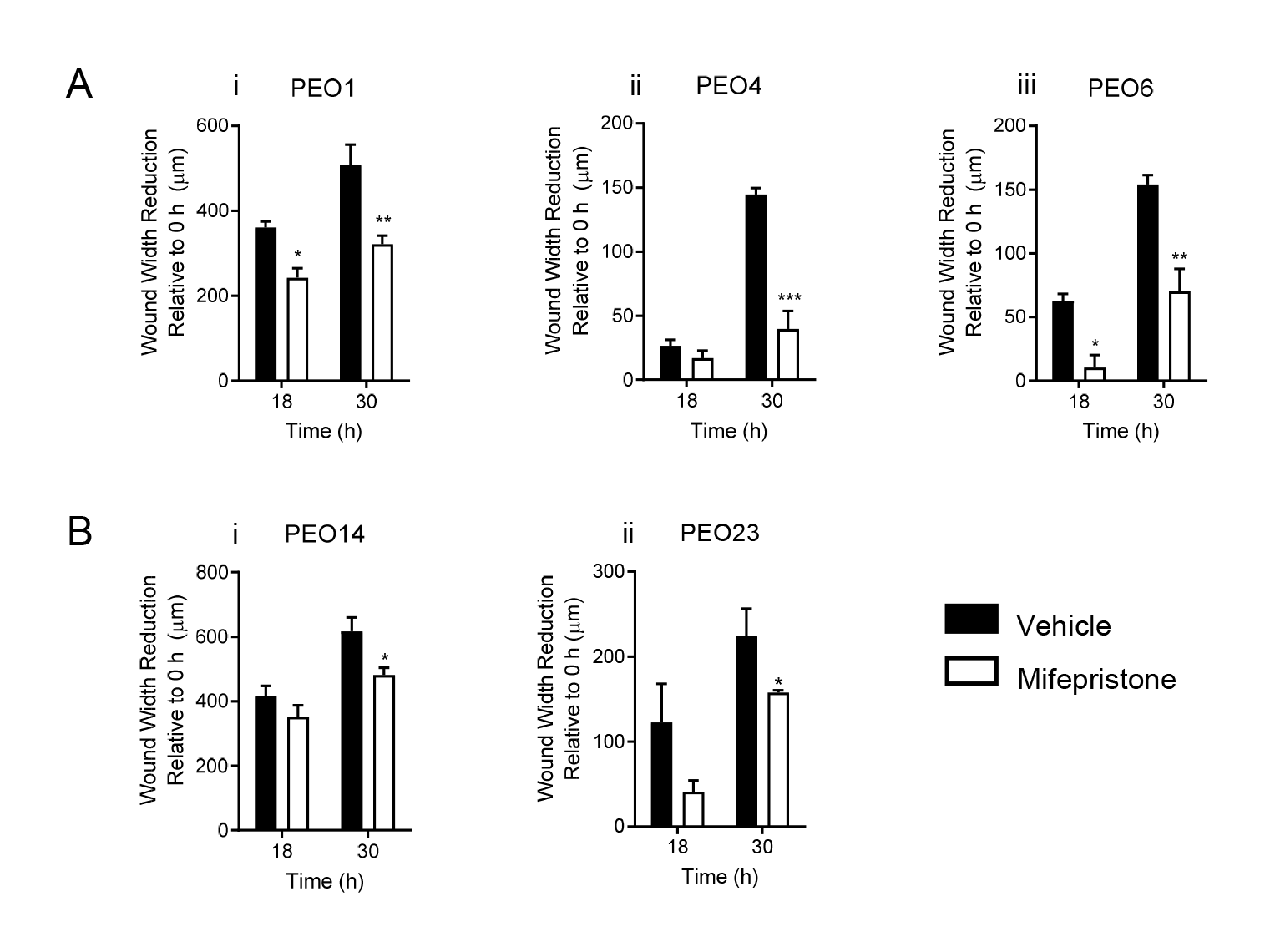
